# Bacterial resistance and co-existence with temperate bacteriophages in the human gut

**DOI:** 10.64898/2025.12.16.694740

**Authors:** Noreen Lanigan, Hugh H. Harris, Neolene Almeida, Colin Hill, Alex R. Hall, Pauline Deirdre Scanlan

**Author notes:** These authors contributed equally to this work (Noreen Lanigan, Hugh H. Harris).

## Abstract

Our understanding of the eco-evolutionary dynamics and consequences of bacteria-phage interactions in nature is limited. By combining longitudinal sampling and analysis of *Enterobacterales* and associated phages isolated from the human gut, with experimental evolution, our work identifies mechanisms that underpin observed patterns of temperate phage-bacteria co-existence and bacterial resistance in this clinically relevant microbiome. Specifically, we show that limited phage infectivity observed against sympatric and allopatric hosts can be explained by the prevalence of spontaneous prophage induction (SPI) bacterial phenotypes and ecological shifts and turnover in focal host populations at multiple taxonomic levels. Despite the ubiquity of temperate phages in the human gut, these findings challenge the assumption that they are major drivers of within-host bacterial evolution and diversification over short-time scales in this natural ecosystem.

## Main Text

Bacteriophages (phages) are predicted to have wide ranging effects on the ecology and evolution of bacterial populations in the human gut (*1–4*). Evidence for phage mediated selection on bacterial hosts in this ecosystem is apparent from the presence of diverse phage defence mechanisms (*5*), together with variations in putative phage receptor genes and prophage regions between strains of different species (*6*). Genomic analysis has showcased the evolution of a novel (lytic) phage species within an individual microbiome (*7*) and highlighted extensive inter- and intra-personal variation in the diversity and stability of phages over time (*8*). Complementary studies using mouse models have demonstrated the importance of ecological context and specific bacteria-phage combination(s) in shaping the dynamics and outcomes of their interactions with respect to key traits such as *de novo* resistance evolution (*9–11*) and lytic phage host range expansion (*11*). *In vivo* models have also highlighted the contribution that temperate phage mediated horizontal gene transfer (HGT) makes to within-host adaptation and colonisation dynamics (*12*), as well as the potential for lytic phages to reduce specific host populations and in turn affect ecological dynamics resulting in alterations to community level functionality (*10*). However, most sequence-based studies are cross-sectional and limited by numerous issues that are well recognised and discussed in detail elsewhere (*13–15*). Furthermore, insights from *in vitro* models, like all laboratory simulations of real-world scenarios, are confounded by the use of allopatric combinations of bacteria and phages whose interactions are often embedded within a relatively low diversity microbial environment. Accordingly, our understanding of bacteria-phage interactions in the human gut is incomplete and perhaps even artefactual. Crucially, there have been no direct empirical tests of the temporal dynamics of bacteria-phage interactions *in natura,* including whether bacteria and phages can co-evolve and drive within-host microbial diversification in the human gut over short, but ecological and evolutionary relevant microbial timescales of weeks and months.

To address this fundamental knowledge gap in our understanding of the human gut microbiome we aimed to capture and study the eco-evolutionary dynamics of naturally occurring, non-model bacteria and phages isolated from this ecosystem. To do so, we sampled seven healthy infants and seven healthy adults at three timepoints over a 12–15-week period. We focused our study on predominant *Enterobacterales* genera and species, such as *E. coli,* and associated phages owing to their ecological and clinical relevance (*16*, *17*), as well as their experimental tractability which facilitated the scale of analysis required. We first isolated 32 bacterial colonies of our focal population from all three faecal samples from each individual, together with a filtrate of each faecal sample containing potentially infective phages. We then used state-of-the-art methodologies that combine cross-inoculation time-shift assays and measures of local adaptation (LA)(*18–20*) of isolated bacteria and phage filtrates with the aim of generating bacteria-phage interaction matrices (*21*) that could identify patterns of phage population level infectivity and bacterial resistance within and between individual gut microbiomes. Each faecal filtrate was also tested against indicator bacterial hosts (*4*) as a control to assess for the presence of viable phages populations that could infect our primary focal bacterial species (*Escherichia coli*). This analysis showed that phages capable of infecting our focal taxon were present in many samples, however, phage infectivity was very rare against bacterial isolates sampled from the gut and only observed in sympatry for two of the 42 different samples obtained from the 14 different individuals tested. Further analysis of phage infectivity at the individual level, using phages isolated from these two samples, revealed that a signal for phage local adaptation in time within these two individuals could in fact be explained by temporal variation in the relative abundance of sensitive hosts that co-existed with diverse and intrinsically resistant bacteria. To complement this analysis, we then co-cultured sympatric bacteria–phage pairs *in vitro* over time. Whilst our co-culture models demonstrated the potential for temperate phages to drive extensive genetic variation within and between bacterial populations, this analysis also highlighted how co-evolutionary dynamics could be constrained by resistance costs and the *de novo* evolution of spontaneous prophage induction (SPI) phenotypes. Notably, these SPI phenotypes produced phages without external induction while remaining resistant to them at the population level. This finding prompted us to test whether SPIs could in fact explain the paradoxical co-existence of phages alongside resistant hosts observed across many of our original faecal filtrate time-shift assays. Using a combination of phenotypic screening, SPI-derived phage host range analyses, and detailed taxonomic and genomic characterisation of bacterial isolates, this analysis revealed that although SPI prevalence accounted for many cases where no infectivity in sympatry was observed, the broader ecological dynamics of diverse, intrinsically resistant host populations also explain the consistently high resistance to phages observed in the human gut microbiome.

## Results

### Phage infectivity is rare with temporal variation in sympatry underpinned by ecological shifts in *Enterobacterales* diversity at multiple taxonomic levels

Viable phages capable of infecting *E. coli* indicator hosts (*4*, *22*) were detected in more than a third of faecal sample filtrates (15/42). However, phage infectivity, at the population level, against sympatric and allopatric bacterial hosts isolated from faecal samples, was very rare (Fig. 1A and 1B). Here, phage infectivity was only observed for 26 of 3136 phage populations *versus* individual bacterial colony isolates (0.8%), with positive cases of infectivity only observed in sympatry and at contemporaneous time-points for individuals I2 and A4 (Fig. 1A-C). Subsequent analysis of phage infectivity at the individual level, using phages isolated from population filtrates tested against sympatric hosts, revealed that phages isolated from I2 phage populations against sympatric bacteria at T1 also had the same infectivity phenotype as each other, and although the infectivity of these phages was consistent with the population level analysis for T1, these phages could infect an additional two bacterial hosts from the earlier time-point of T0 and 18 hosts from the later time-point of T2 (Fig. 1E). Similar analysis revealed that individual phages isolated on sympatric bacteria from A4 at T0 had the same infectivity profile as each other and corresponded exactly with phage population level analysis when tested against sympatric bacteria from T0, T1 and T2 (Fig. 1H). {Citation}

**Fig. 1.**
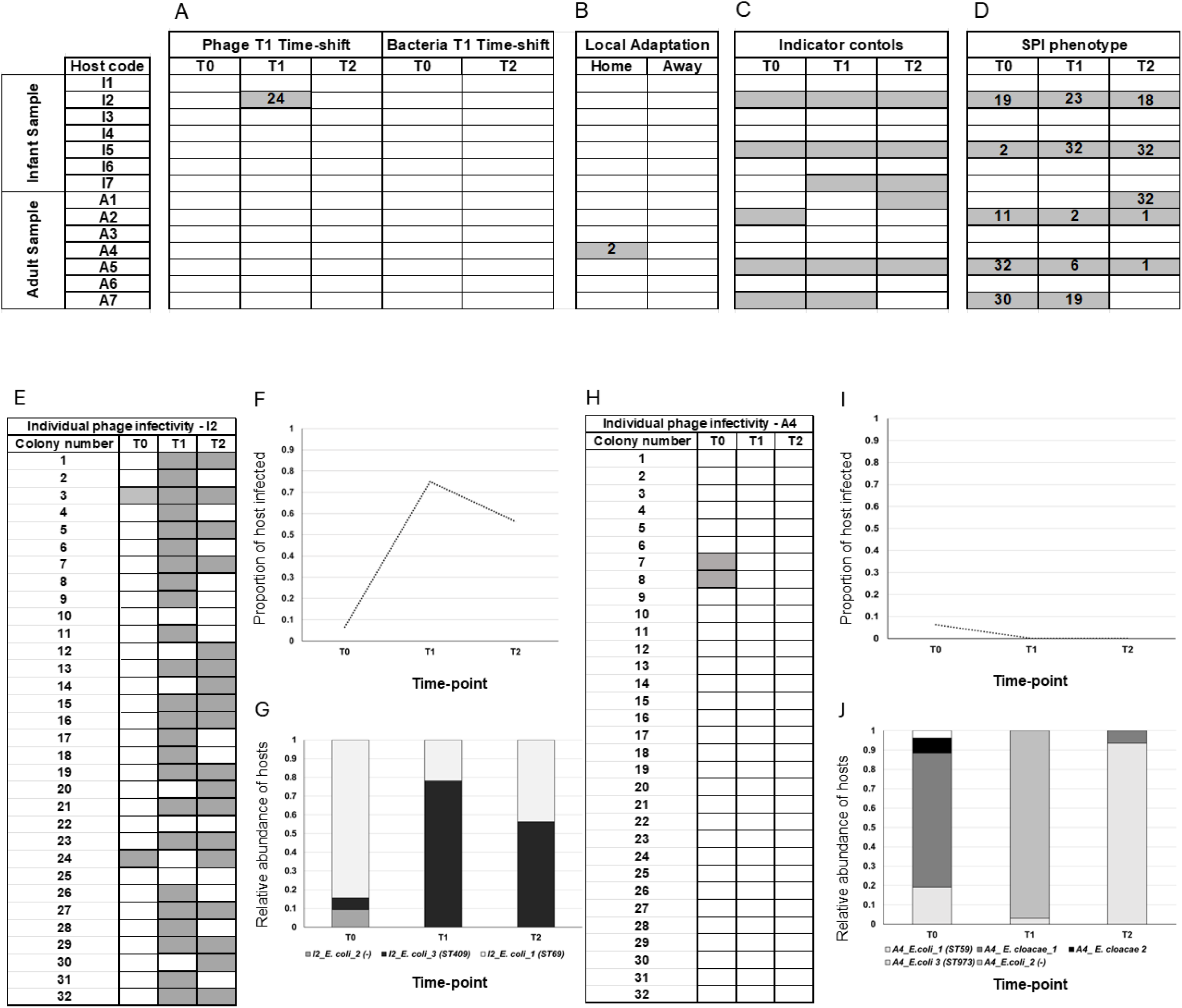
Summary of phage infectivity at population and individual levels across sympatric, allopatric, and indicator hosts, together with survey of SPI phenotype prevalence across from individual hosts. **(A)** *Time-shift assays:* Phage T1 time-shift – phage populations sampled from T1 were tested for infectivity against 96 bacterial isolates from that same individual, 32 from each time-point (T0, T1, and T2); Bacterial T1 time-shift - bacterial isolates from T0 and T2 (32 from each timepoint) were tested for sensitivity to T1 phage population infectivity: white squares indicate no infectivity across all 32 isolates for that individual and time-point; grey squares show the number of host isolates (out of 32) sensitive at the population level for that individual and timepoint. **(B)** *Local adaptation assays* - Phage population filtrates from one individual at a given time point were tested against the 32 bacterial isolates from the same individual/time-point (home/sympatry) and against 32 isolates from a different individual (away/allopatry): white squares indicate no infectivity; grey squares show the number of host isolates (out of 32) sensitive at the population level for that individual and timepoint. **(C)** Phage population-level infectivity from all samples against indicator hosts (binary assay white = no infectivity against indicator hosts, grey = phage plaques present on indicator hosts). **(D)** Prevalence of SPI phenotypes per individual sample and timepoint (white = no SPIs present out of 32 hosts tests; grey = SPI present in sample with the number of hosts, of 32 tested per sample, with this phenotype. **(E-J)** Individual-level infectivity of ph_I2_188 (isolated in sympatry from sample T1 of individual I2) and ph_A4_364 (isolated in sympatry from sample T0 of individual A4) against 96 bacterial isolates sampled across T0, T1, and T2 respectively shown as binary matrices (white = no infectivity, grey = infectivity) with corresponding temporal infectivity plots **(F, I) and s**train-level relative abundance profiles **(G and J)** of sensitive and resistant bacterial hosts over time for individuals I2 (*E. coli*_ST409 = sensitive host population) and A4 (A4*_E. cloacae*_2 = sensitive host population).

Whilst these patterns of phage infectivity against sympatric hosts could superficially be interpreted as evidence for phage local adaptation in time (Fig. 1F and I), an alternative explanation is that temporal variation in phage infectivity could be due to species and strain level bacterial diversity within samples and/or ecological shifts in bacterial composition within these two individuals over time. In other words, this could represent an ecological signature of phage-bacteria interactions, rather than co-evolution within interacting populations (*23*). To test these contrasting hypotheses, we used comparative genomic analysis of sensitive and resistant bacterial hosts isolated from all three-time points for individuals I2 and A4. For individual I2, genome resolved analysis of isolates sampled over time separated them into three *E. coli* sequence types (STs) (Fig. 1G, fig. S1). Isolates of only one of the three STs was sensitive to the phages tested (*E. coli* ST409), and the relative abundance of this phage sensitive ST at each time point mapped over the infectivity profile of individual phages observed in our time-shift analysis (Fig. 1F). Similarly, for individual A4, both phage-sensitive hosts from T0 were identified as a strain of *Enterobacter cloacae* (designated A4_*E. cloacae_*2 for disambiguation and referencing). This strain was absent at T1 and T2 where no phage infectivity was observed (Fig. 1H and J) and the remaining resistant T0 isolates, as well as resistant hosts from T1 and T2, could be disambiguated into different genera, species and strains of *Enterobacterales* highlighting considerable ecological turnover in the relative abundance of different bacterial populations at multiple taxonomic levels within individual A4 over time (Fig. 1J, fig. S2). Collectively, genome-resolved analysis of potential host populations for individual I2 and A4 show that patterns of resistance and sensitivity to phages in the gut can be explained by genus, species, and strain level ecological shifts in the relative abundance of sensitive hosts within and between time-points rather than rapid cycles of resistance and infectivity evolution within a bacterial population giving rise to a signal for phage local adaptation in time.

### Temperate phages and bacteria can co-exist over time, but cycles of antagonistic coevolution are constrained

To better understand the temporal dynamics and genetic basis of resistance evolution to temperate phages we established two independent co-culture *in vitro* models using sympatric bacteria-phage pair isolates from I2 and A4; an *E. coli* strain from I2 (I2_T1_19) together with its infective phage (ph_I2_188) and an *Enterobacter cloacae* strain from individual A4 (A4_T0_8) and its infective phage (ph_A4_364). Both phages were sequenced and identified as novel temperate phages with no known taxonomic affiliation (fig. S3, table S1). Whilst bacteria and phage populations co-existed at high population sizes over time *in vitro* (10 days, ∼70 bacterial generations; Fig. 2A and B), tests for signals of bacteria-phage coevolution using time-shift assays provided no evidence for directional changes in phage infectivity over time (slope of infectivity vs phage time, estimated separately for all 16 bacterial colony isolates in each population, *p* > 0.05 in one-sample *t*-tests against a mean of zero for each population; Fig. 2C and D; fig. S4). This was true for both phage-bacterium combinations tested and additional experiments with bacterial isolates sampled from the end (T10) rather than the middle (T5) for both co-culture models supported a similar picture (fig. S5). On the bacterial side, we detected increasing resistance over time, but only in the A4_T0_8 versus ph_A4_364 combination (mean slope of infectivity across bacterial time, estimated for each replicate population and then tested against a mean of zero by one-sample *t*-test: *p*=0.005; Fig. 2E) with the majority of bacterial isolates at T10 in each population resistant to phages from T5. For the other combination (I2_T1_19 and ph_I2_188), we did not detect directional changes in phage infectivity (mean slope of infectivity vs bacteria time not different from zero: *p*>0.05), with the proportion of hosts infected instead lowest against bacteria from the same time point as the tested phage lysate, e.g. T5 (Fig. 2F). However, this trend was not sustained to the end of the experiment with phage lysates from transfer 10 almost always infectious against bacterial isolates from transfer 10 in this phage-bacteria combination (fig. S5). Moreover, bacterial colonies from T10 in all populations were also sensitive to the ancestral phage and phage from T5, indicating limited resistance evolution in this co-culture model pairing (fig. S5). Collectively, these data show that phage infectivity did not evolve directionally in either phage-bacterium combination, and although we detected resistant bacteria in both, the outcome differed between the two different bacteria-phage sympatric combinations. Notably, bacterial isolates in both combinations often had the same resistance/sensitivity phenotype against phages from all tested timepoints (fig. S4 and S5). This data, together with the finding that there appears to be little to no change in the phage population-level infectivity phenotype relative to the ancestral phage used to inoculate the experimental model, indicates limited phage infectivity evolution over the timescale of these experiments. As such, rapid cycles of antagonistic co-evolution, that are commonly observed in *in vitro* models using lytic phages (*24–27*), and more recently in natural populations (*28–30*), appear constrained between pairs of sympatric bacteria and temperate phages isolated from the human gut.

**Fig. 2.**
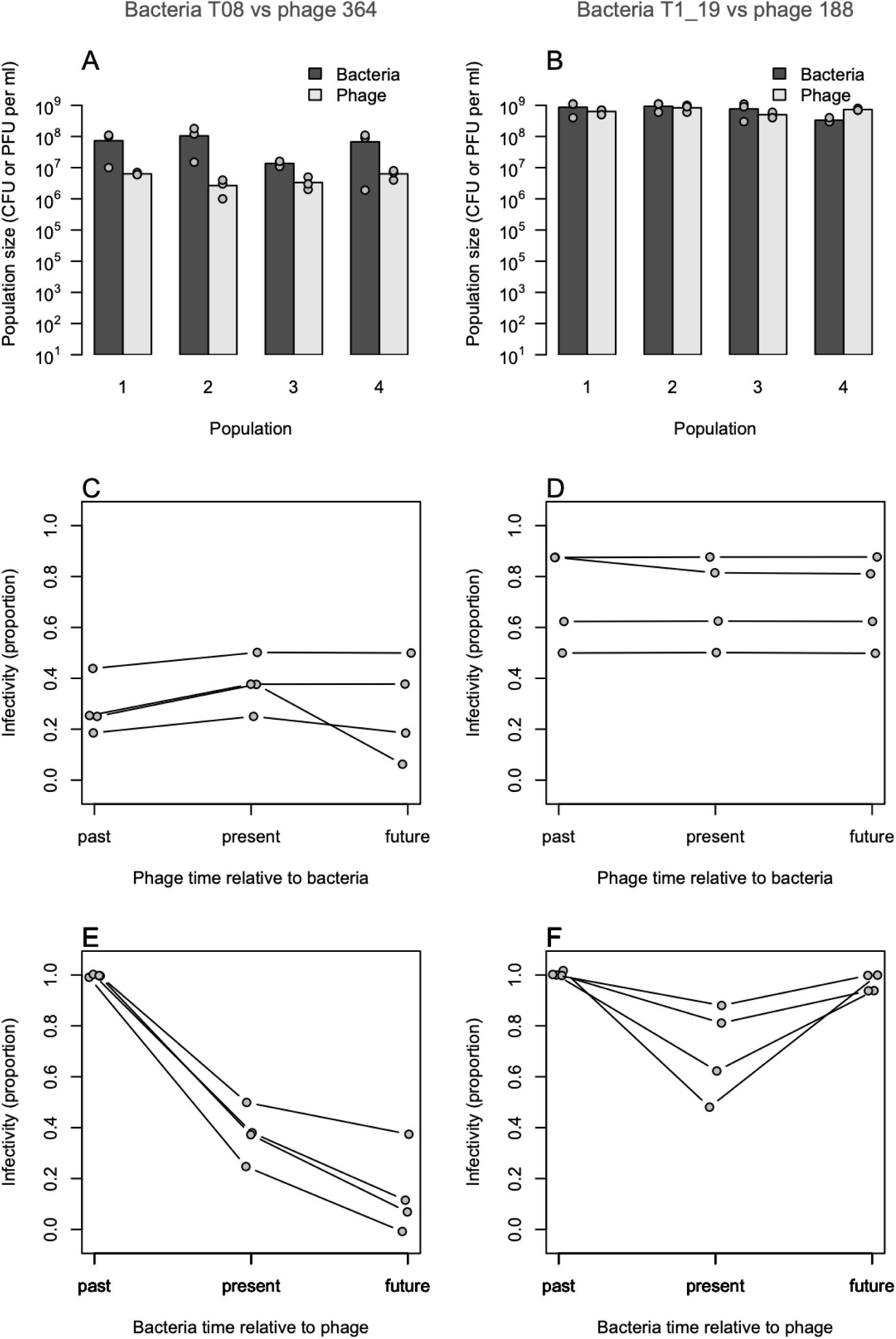
Bacteria-phage co-existence and co-evolutionary interactions *in vitro*. **(A and B)** Quantitative analysis of bacteria and phage population sizes showing mean CFUs or PFUs/ml ± 1.s.d per replicate population) at the last experimental time-point (T10) showcasing the ability of phages to co-exist with their bacterial hosts over time in our co-culture experiments; **(A)** Count data from A4_T0_8 and ph_A4_364 co-culture model and **(B)** count data from I2_ T1_19 and ph_I2_188 co-culture model. **(C-F)** shows evolutionary interactions between bacteria and phages during co-cultivation. For each of two bacteria-phage combinations (columns: A, A4_T0_8 and ph_A4_364, and B, I2_ T1_19), the upper panels show infectivity of phage lysates from transfer 0 (past), 5 (present), or 10 (future) of the co-culture experiment when tested against 16 bacterial colony isolates from transfer 5. Each point shows the proportion of the tested isolates that could be infected, and each series (points and lines) shows this for one of four replicate populations. The lower panels show infectivity of phage lysates from transfer 5 when tested against the ancestral bacterial clone (past), 16 bacterial colony isolates from transfer 5 (present), or 16 independent colony isolates from transfer 10 (future).

### Temperate phages drive the genetic diversification of bacterial populations *in vitro* but resistant hosts *in natura* lack phage receptors required for infectivity

To identify the genetic basis of resistance to temperate phages evolved *in vitro* we performed comparative genomic analysis of wild-type I2_T1_19 and A4_T0_8 and their evolved phage resistance mutants from our *in vitro* models. This analysis demonstrated that temperate phages can drive the rapid genetic diversification of hosts both within and between populations, with both extra-cellular and intra-cellular mechanisms of resistance and immunity evolving *de novo* (Fig. 3A, table S2). A subset of I2_T1_19 mutants evolved resistance *via* lysogenisation with ph_I2_188 either integrated into the host chromosome or present as an episomal circular contig. Analysis of the remaining I2_T1_19 mutants showcased diverse molecular mechanisms underpinning receptor mediated resistance evolution (table S2). In two resistant mutants, that evolved in independent populations, intragenomic recombination between two homologues of a phosphomannomutase (*pmm*) gene flanking the colanic acid biosynthesis operon resulted in the formation of a single *pmm* hybrid gene and the deletion of 15 contiguous genes (totalling 24.54 kb or 0.5% percent of the bacterial genome) (Fig. 4A, table S3). Whilst colanic acid capsular polysaccharide overproduction can underpin generalist phage resistance strategies in laboratory strains of *E. coli* (*31*) it is also a recognised phage receptor for *Enterobacterales* species (*32*). This large-scale deletion might also be linked to phage resistance *via* its effects on O-antigen biosynthesis given the overlap between both biosynthetic pathways (*33*, *34*). This hypothesis is supported by analysis of the remaining I2_T1_19 resistant mutants which had mutations in genes that formed part of the LPS operon. Notably these mutations were mediated by the insertion of different IS elements, namely IS5 and IS1 InsB, into the O-antigen ligase in two of the mutants (IS5), and into a glycosyltransferase gene (IS1 InsB) in the other mutant (Fig 3A, table S2). Similar to I2_T1_19 mutants, a subset of A4 mutants evolved immunity *via* lysogensiation with ph_A4_364. The remaining mutants evolved resistance *via* receptor modification with SNPs and indels resulting in premature stop codons in genes encoding for O-antigen ligase and glycosyltransferases (Fig 3B, table S2). This further highlights the importance of LPS as a phage receptor in clinically relevant gut species and how genetic variation in the genes encoding this trait underpins resistant phenotypes.

**Figure 3.**
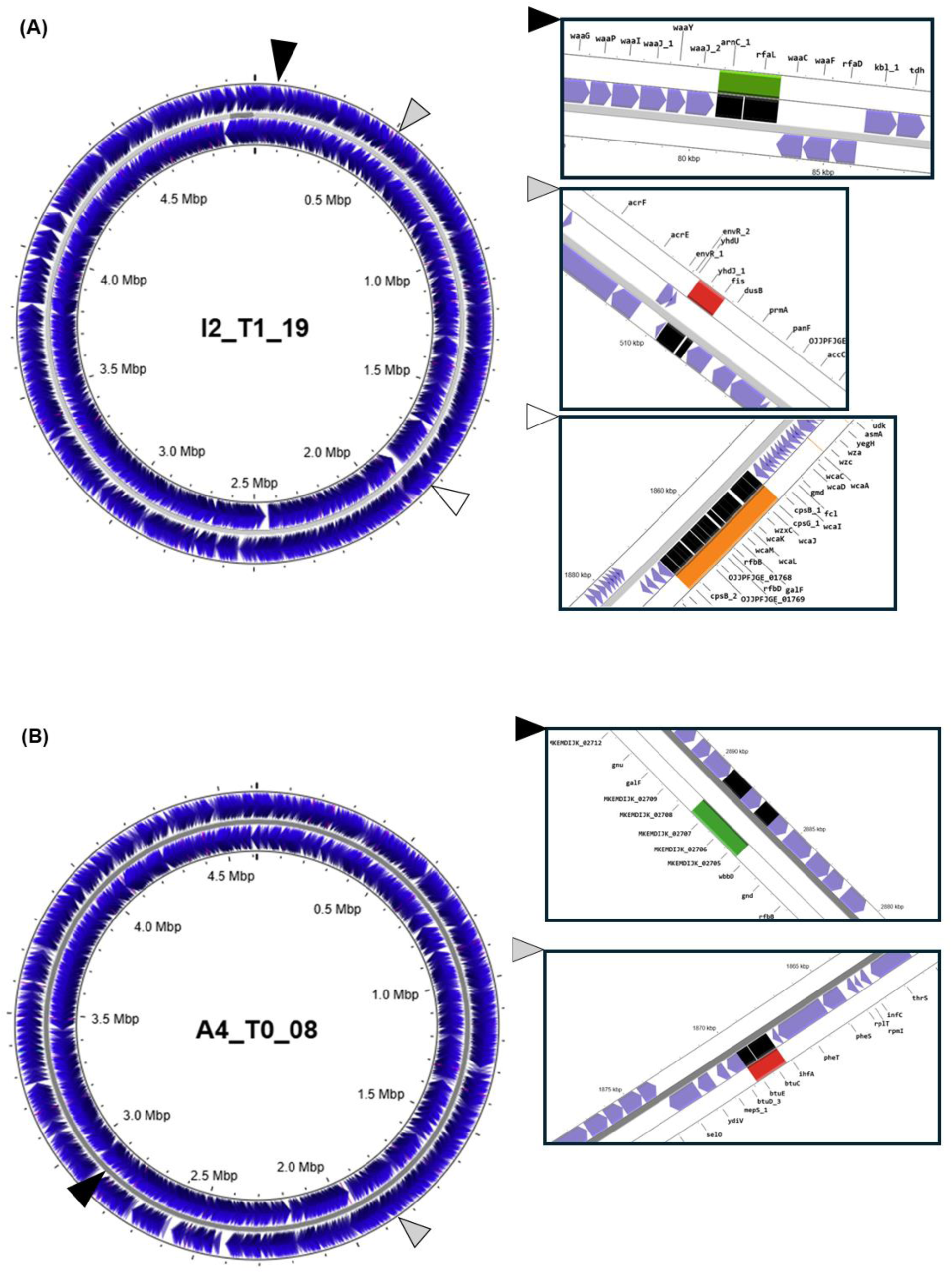
Genomic regions and mutational events underpinning host resistance evolution to phages. **(A)** Circular genome of *Escherichia coli* strain I2_T1_19 showing the loci involved in resistance evolution to ph_I2_188 for evolved mutants in co-culture experiments. The outer and inner genomic rings display all genes on the forward and reverse strands of the ancestral genome, respectively. The black, grey and white triangles denote, respectively; the insertion sites for transposable element insertion into glycosyltransferase (arnC_1) or an O-antigen ligase (rfaL) genes, the integration site of phage_188 between genes yhdJ_1 and fis, and the 15-gene deletion between cpsG_1 and cpsG_2 involving homologous recombination of flanking phosphomannomutase genes. **(B)** Circular genome of *Enterobacter cloacae* strain A4_T0_08 showing loci involved in resistance evolution to ph_A4_364 for evolved mutants in co-culture experiments. The outer and inner genomic rings display all genes on the forward and reverse strands, respectively. The black and grey triangles denote, respectively insertion-deletion events in genes encoding a glycosyltransferase (locus tag MKEMDIJK_02705) or an O-antigen ligase (locus tag MKEMDIJK_02707) which conferred phage resistance, and the phage integration site of phage ph_364 between genes btuC and btuE.

**Figure 4.**
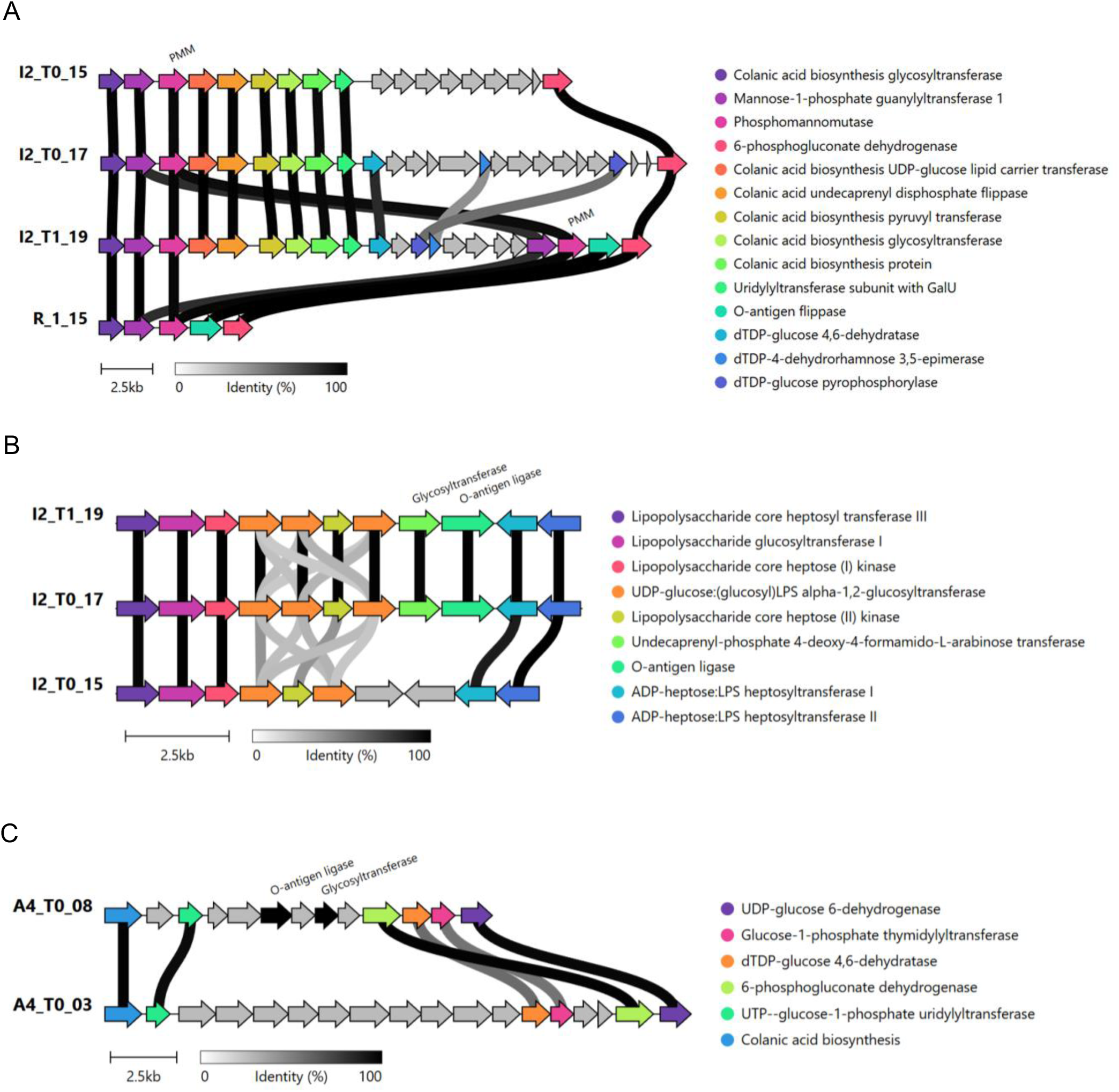
Comparative genomic analysis of PMM and LPS operons underpinning receptor mediated phage resistance evolution. **(A)** Comparative genomic analysis of PMM operon in representatives of naturally co-occuring phage sensitive (I2_T1_19) and phage resistant *E. coli* strains temporally sampled from the gut microbiome of individual I2 together with an evolved resistant mutant of T1_19 (namely, R.1.15) from *in vitro* co-culture experiments highlighting both gene presence/absence and percentage similarity in natural isolates, and an intragenomic homologous recombination event resulting in a 24.5kb deletion in evolved resistant mutant R.1.15. **(B)** Comparative genomic analysis of LPS operon encoded by representatives of naturally occurring phage sensitive (T1_19) and phage resistant *E. coli* strains temporally sampled from I2 showing genetic variation in lipopolysaccharide operon and **(C)** comparative genomic analysis of LPS operon of phage sensitive and resistance variants of *Enterobacter cloacae* natural isolates from individual A4 showing genetic variation in lipopolysaccharide operon.

Whilst MLST, GDBP, and ANI analyses showed that the co-occurring phage-sensitive and phage-resistant isolates of *E. coli* isolated from individual I2 and *Enterobacter cloacae* isolated from individual A4 were phylogenetically distinct strains (fig S1 and S2), the underlying genetic basis of resistance or immunity between strains of these species was not known. Using our *in vitro* genomic data as a guide we first ruled out specific prophage-mediated superinfection exclusion (SIE) as the mechanism given that naturally co-occurring resistant isolates of both strains sampled from the gut microbiomes of I2 and A4 did not encode their respective prophages (e.g. ph_I2_188 or ph_A4_364). Further comparative genomics of *E. coli* isolates from individual I2 revealed that all phage-sensitive *E. coli* sequenced encoded intact and genetically identical PMM and LPS operons. However, co-occurring resistant *E. coli* strains from individual I2 lacked several genes in the PMM operon and/or LPS pathway (Fig. 4A and B: table S3 and S4. As these traits underpin resistance evolution, the absence of these specific genes and divergence within the PMM and LPS operons demonstrates that co-occurring (sympatric) resistant *E. coli* strains in the human gut lacked the specific receptor structure required by phage ph_I2_188 to infect hosts. Similarly, comparative genomic analysis of phage-sensitive isolate A4_T0_8 with resistant co-occurring *E. cloacae* strains revelated that key LPS biosynthesis genes (e.g., glycosyltransferase, O-antigen ligase) required for infection by ph_A4_364 were also missing and genetically divergent (Fig. 4C; table S5). Taken together with earlier evidence showing how strain-, species-, and genus level diversity within individuals underpinned the bacteria-phage infectivity matrix (Fig.1 E-J), these results demonstrate that the genetic basis of intrinsic resistance in co-occurring strains of the same species in the gut is underpinned by the absence of and genetic variation in receptor encoded traits.

### Spontaneous prophage induction phenotypes facilitate bacteria-phage co-existence and confer cost-free or low-cost population-level immunity

Recent work has demonstrated that spontaneous prophage induction (SPI) phenotypes are a common consequence of temperate phage infection and host lysogenisation (*35*). A key aspect of this phenotype is that a subset of the population can undergo spontaneous lysis and release phages into their local environment, but these SPI derived phages cannot infect remaining individuals in the population owing to superinfection exclusion (SIE) (*35*, *36*). We hypothesised that these specific phenotypic traits of SPI bacteria might help explain the co-existence of temperate phages with limited infectivity against bacterial hosts *in vitro*, particularly in our A4 model. To test this, we spotted filtered supernatants of overnight cultures for all A4_T0_8 and I2_T1_19 lysogenized resistant mutants against their ancestral hosts. This assay demonstrated that all mutant lysogens had an SPI phenotype and produced phages (A4_T0_8 SPI mutants; mean 1.8 x 10^8^ PFUs/ml; range 1.0 to 3.2 x 10^8^ PFUs/ml, I2_T1_19; mean 1.1 x 10^7^ PFUs/ml; range 4 x 10^6^ to 2.2 x 10^8^) that could infect their respective ancestral host but could not infect their lysogenised mutant host populations. This data supports our prediction that the apparently paradoxical co-existence of phages with resistant (immune) hosts *in vitro* can be explained in part by the prevalence of SPIs, particularly in the A4_T0_8 model.

Although SPIs and resistance did evolve in our I2 co-culture model it was relatively rare compared to the A4 co-culture model with one possible explanation being that costs associated with resistance evolution differed between the models. Consistent with this prediction, both mutant classes (SPI/lysogens and receptor mutants) in our A4 model had slightly higher average ODmax compared to the ancestor, and although not statistically significant, this slightly higher ODmax for SPI/lysogen (two-tailed one sample t-test; t = 1.9, df = 4, p = 0.13) or receptor mutants (two-tailed one sample t-test t = 0.829, df = 2, p = 0.494), (Fig. 5A), helps explain the rise and maintenance of resistance for this bacteria-phage pair model *in vitro*. Analysis of I2_T1_19 phage resistant mutants revealed that lysogens had lower average ODmax readings compared to the WT (two-tailed one sample t-test; t= -2.09, df 3, p = 0.128) and receptor mutants had a significantly lower ODmax compared to both the WT (two-tailed one sample t-test; t = - 3.54, df 4, p = 0.024) and lysogens (two-tailed unpaired t-test; t = 2.55, df 7, p = 0.038) (Fig. 5B) which helps explain selection against phage resistance for host I2_T1_19 when co-cultured with ph_I2_188 *in vitro.* Crucially, this data demonstrates how selection against costs of resistance, as well as the type of resistance or immunity evolved, can constrain cycles of co-evolution by either removing the selection pressure to evolve counter-infectivity in the phage population in the case of high cost resistance, or by potentially shifting the selection pressure from hosts onto phage populations that compete for hosts as resources. In the latter scenario, it is possible that the continual release of phages by SPI phenotypes may still facilitate evolutionary innovation and diversification over the longer term by driving competition between temperate (extracellular) phages and prophages for bacterial hosts, for example by selecting for escape defence mutants (*37*). This idea is consistent with recent evidence highlighting how different prophage encoded defence strategies protect their hosts (their sole resource) against infection by other temperate coliphages (*38*) and highlights the increasingly recognised and important role of phage:prophage competition as a driver of bacterial diversification over time. Moreover, the *de novo* evolution of SPI phenotypes in response to sympatric phage predation adds to recent studies demonstrating the importance of host phenotypic variation i.e. host physiology (*39*) and phase variation (*40*) in sustaining bacteria–phage coexistence, and in doing so, highlights how non-genetic factors can also affect the ecological and evolutionary dynamics and outcomes of bacteria-phage interactions in nature.

**Figure 5.**
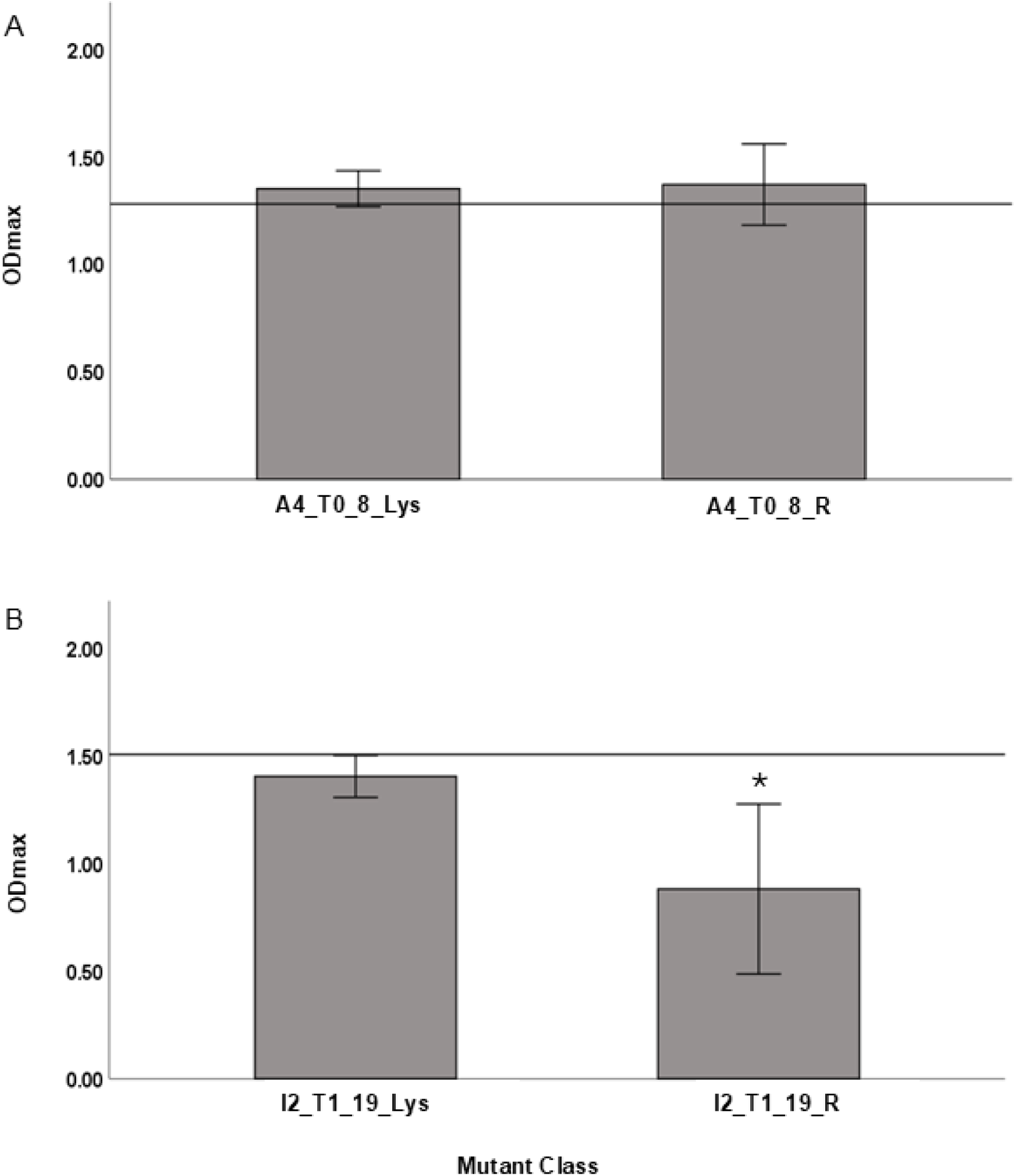
Costs of resistance to temperate phage vary by bacteria-phage combination and type of resistance evolved. Costs of resistance for each bacteria-phage combination and vased on genetic basis of resistance evolution (mutant class). ODmax analysis of lysogenised (L) and receptor (R) resistance mutants for **(A)** A4_T0_8 and ph_A4_364 bacteria-phage pair and **(B)** I2_T1_19 and ph_I2_188 bacteria-phage pair. Straight line represents the ODmax of each respective WT ancestor with * denoting a significant difference in OD max relative to WT (*P* < 0.05).

### SPIs phenotypes are common in the human gut and help explain the observed co-existence of phages with resistant hosts

The rapid selection for SPI phenotypes in our co-culture experiments helps to explain the co-existence of resistant bacteria with high numbers of phages *in vitro*. We further hypothesised that the presence of SPI phenotypes in the gut microbiomes of the different individuals sampled could explain the co-existence of phages with resistant bacteria *in natura*, as evidenced by the paradoxical rarity of phage infectivity against sympatric hosts but the presence of viable infective phages against indicator hosts *E. coli* MAC and C observed in the original population time-shift and control assays (Fig. 1A-C). To test this hypothesis, we screened all 96 bacterial isolates sampled from each infant (n = 3) and adult (n = 4) whose faecal filtrate(s) were positive for phage infectivity against *E. coli* MAC and C in the initial time-shift assays (Fig. 1C) for potential SPI phenotypes (n = 672 bacterial isolates tested). SPI phenotypes were present to varying degrees in all four adults and two of the three infants that were positive for phage infectivity in the original assay (Fig.1C and D). In the third infant (I7) that was positive for phages at T1 and T2 against MAC and C, no SPI phenotypes were detected for any of the 96 bacterial isolates. This indicates that the phage(s) detected on MAC and C for infant I7 were non-SPI derived or derived from a bacterial host that was not present within the subset of bacteria sampled. Aside from this case, the detection of phages against MAC and C in our initial phage population level screen is consistent with the screening of potential SPI positive samples in both infants and adults (Fig. 1C and D). Representatives of each unique bacterial strain type (n = 49) identified through strain-typing PCRs of all 96 bacterial isolates from each of the seven remaining individuals that were negative for phage infectivity against MAC and C (total 96*7 = 672 isolates) were also tested for an SPI phenotype. Only one additional SPI strain type was detected (from individual I6), which further supports our hypothesis that phages captured in the original phage population screen against MAC and C were likely SPI derived given the consistency of the results of that assay and the screening of individual bacterial isolates from positive and negative samples against MAC and C. Next, we tested filtered supernatants derived from SPI bacterial overnight cultures of each unique SPI strain type for infectivity against all sympatric bacteria from that same individual. Consistent with the superinfection exclusion observed for our *in vitro* evolved phage resistant SPI mutants we found that none of the SPI derived phages could infect isolates of the host strain type from whence the phages were derived. Moreover, no SPI derived phage infectivity was observed against any other (non-SPI) bacterial strain types present within the same individual, with one exception for individual A1. Here, two different *E. coli* STs (ST69 and ST4377) present at T2 had an SPI phenotype, but only phages derived from *E. coli* ST69 bacteria could infect a different *E. coli* strain (ST646) present in this individual at the earlier timepoints of T0 and T1.

### Temporal ecological dynamics of bacterial hosts at multiple taxonomic levels and phage specificity also account for patterns of resistance to temperate phages

Whilst the presence of SPI-positive bacteria helps to explain phage co-existence with resistant hosts in some individuals, in many cases no infective phages were detected against *E. coli* indicator hosts MAC and C in the faecal filtrate phage screen (Fig. 1C), and non-SPI strains were present in many samples (Fig. 1D). Based on patterns of phage infectivity data observed for individuals I2 and A4 (Fig. 1E-J), we hypothesised that broader bacterial host diversity within and between individuals i.e. the presence of taxonomically diverse and intrinsically resistant (nonhost resistance (*41*)) bacterial populations might also help explain the absence of infective phages against *E. coli* indicators in some samples.

To analyse both the taxonomic diversity and compositional turnover of bacterial hosts within individuals in this study we first analysed PCR-based strain typing for all 1,344 bacterial isolates sampled across time from individuals in this study and assigned taxonomy to unique strain isolates using 16S rRNA sequence analysis. Five of the seven individuals that were negative for infective phages in faecal filtrates and SPI phenotypes when tested against *E. coli* indicator hosts were primarily colonised by non-*E. coli* taxa, including *Enterobacter, Citrobacter, Klebsiella,* and *Enterococcus* (fig. S6). Within the infant cohort, I1, I4, and I6 were dominated by non-*E. coli* strains. Similarly, two of three adults (A4 and A6) were also colonised by non-*E. coli* taxa, with A6 stably harbouring *E. marmotae* across all timepoints (fig. S6). Further Bray–Curtis dissimilarity analysis of bacterial diversity data demonstrated that while some individuals were stably colonised by the same *Enterobacterales* strain(s) over time, many showed rapid shift and ecological turnover in bacterial composition across genera, species, and strains with the predominant *Enterobacterales* diversity often shifting completely between timepoints (mean Bray–Curtis dissimilarity between T0 and T2 = 0.46, range 0–1; fig. S7). Given the typically narrow and strain-specific host range of phages (4, 38), the absence of host *E. coli* populations within some samples helps explain the lack of infective phages, including those that are SPI derived, detected against *E. coli* indicators in our original screen (Fig. 1C). These rapid shifts in the composition of bacterial populations within some individuals, also underscores the importance of host ecological dynamics to our understanding of the observed spatio-temporal patterns and outcomes of bacteria-phage interactions in the gut over short-time scales. Crucially, rapid taxonomic shifts in the composition of host populations not only minimises the ecological effects of phages on bacterial populations but ecological turnover is likely an important constraint on phage-bacteria co-evolution in the gut by limiting the continuous supply of susceptible host populations required to sustain reciprocal cycles of co-evolution.

Independent of bacterial taxonomic diversity, the inability to recover phages infective against sympatric or allopatric hosts is striking. However, it is consistent with other studies that demonstrate the rarity of isolating infective phages using diverse hosts from human faeces as bait (*22*, *42*). The rarity of recovering infective phages in sympatry or allopatry could simply be due to the absence of phages capable of infecting specific taxa (which is not testable) or phages present in samples with very narrow host ranges and limited infectivity above the strain level as evidenced for our earlier analysis of I2 and A4 data. To explore this latter idea further, we assessed the infectivity of ph_I2_188, ph_A4_364, six SPI-derived temperate phages together with the lytic model phages T4, T5, T6, T7, and PPO1 against a phylogenetically diverse panel of whole genome sequenced isolates (n = 38) representing the diversity of *E. coli* strains and taxa recovered across hosts sampled in our study. At biologically realistic titres (10⁵ PFU/mL, reflecting average SPI titres from natural isolates), no infections were observed. Plaques were observed against a small number of *E. coli* hosts, as well as *E. marmotae* but only at artificially high titres (>10⁸ PFU/mL) and were often turbid (table S6). These data support similar findings for temperate coliphages shown elsewhere (*4*) and highlights both the extreme specificity of phage infectivity and limited range of susceptible host populations within a single species such as *E. coli* at biologically realistic values.

In this respect the genetic basis of resistance or immunity could be an intra-cellular antiphage defence system. However, studies have shown that mechanisms such as CRISPR are rare in the gut (*43*), with the primary determinant and predictor of phage infectivity for many species including *E. coli* linked to receptors and O-antigen variation (*22*, *44*, *45*).This is consistent with our own analysis of resistance linked to genetic variation in receptor traits for I2 and A4 and additional ECtyper analysis of our strain collection confirmed extensive O- and H-antigen diversity between strains (table S8). These data together with the narrow host range of temperate coliphages shown here and elsewhere (*4*), indicate that in addition to SPI phenotypes, the ecological dynamics and diversity of *Enterobacterales* that encode extensive genetic variation in phage receptor related traits i.e. the presence of nonhost resistant populations (*41*) within and between individuals also helps explain the rarity of successful phage infections and the observed spatio-temporal of phage resistance patterns in the human gut microbiome.

## Discussion

*In vitro* models and studies targeting specific bacterial taxa and their associated phages in natural environments have shown that phages can drive cycles of antagonistic co-evolution and rapid bacterial diversification over weeks to months (*29, 45*, *46*). Contrary to these findings and our own predictions (1), we conclude that temporal variation in phage infectivity against sympatric hosts in the human gut is best explained by ecological turnover in host community composition at the genus, species, and strain levels, rather than by rapid within-host co-evolution of bacteria and phages. More broadly, phage infectivity against sympatric and allopatric hosts was exceedingly rare, despite the frequent recovery of viable phages capable of infecting our focal taxon in control assays. This apparent paradox, taken together with the overall limited phage infectivity observed in time and space, can be explained by two complementary mechanisms: (i) the widespread presence of spontaneous prophage induction (SPI) phenotypes and (ii) intrinsic or nonhost resistance arising from compositional variation and spatial and temporal shifts in bacterial populations across taxonomic scales.

Our *in vitro* models demonstrate the capacity for temperate phages to rapidly select for SPI phenotypes which promote bacteria-phage co-existence and highlight the importance of phage lifestyle and fitness trade-offs in shaping bacterial evolutionary trajectories. Importantly, the lack of detectable fitness costs associated with SPI phenotypes likely explains their prevalence in nature, as shown here and elsewhere (*46*, *47*). Despite the limited understanding of temperate phage ecology that currently exists (*48*, *49*), the widespread occurrence of SPI phenotypes offers a novel framework for interpreting phageome variation in environmental samples. For example, if SPI phenotypes are a common feature of different taxonomic groups in the gut and are common in other ecosystems of interest their prevalence challenges the assumption that external cues are primarily responsible for temperate phage induction and cautions against over-interpreting environmental variation across hosts or habitats as the primary factor underlying viral dissemination and community-level compositional changes. More importantly, if the host populations from whence temperate phages are derived are also immune to them, extra-cellular temperate phages, with narrow host ranges, are unlikely to have any broad-scale ecological or evolutionary top-down effects on bacterial populations in the short-term owing to their limited infectivity. These insights largely apply to temperate phages, which dominate gut viromes, and whether strictly lytic phages exert stronger top-down control and drive alternative evolutionary outcomes remains unknown. However, it is notable that no lytic phages were recovered in our study against sympatric hosts which is consistent with the inability to recover both temperate and lytic phages from gut samples more generally (*22*, *42*).

Collectively, our findings challenge the view that phages are major drivers of within-host bacterial evolution and diversification in the human gut over short-time scales. Instead, we show that short-term patterns of bacteria-phage co-existence, infectivity and resistance observed are shaped more by ecology than by evolution, and are underpinned by phage lifestyle, the genetic basis and costs associated with resistance, together with the dynamics and phenotypic traits of host populations. These findings have broad implications for our understanding of the ecology and evolution of microbial populations in nature and for how bacteria-phage spatio-temporal interactions networks are shaped in environments that are predominated by temperate phages and have high diversity and ecological turnover in host populations.

## Materials and Methods

### Study participants and sample collection

Seven healthy infants (labelled I1-I7, mean age: 19 months; range 10.5 - 30 months) and seven healthy adults (labelled A1_A7: mean age: 37.3 years; range 27 - 59 years) were enrolled in the study. Ethical permission was sought and granted from Cork Research Ethical Committee, protocol numbers APC079 and APC065. Primary exclusion criteria were the use of antibiotics in the three months prior to sample collection. Informed consent was given for sample collection, and each participant provided three samples that were taken between 4 and 6 weeks apart (42 samples in total). Faecal samples were placed at four degrees and frozen at -80 °C upon collection.

### Bacterial isolation and phage filtrate preparation

*Enterobacterales* were isolated from each sample by selective plating on Luria-Bertani (LB) agar plates and incubated aerobically for 24 hours. 32 individual colonies were picked from each of the three faecal samples (i.e. T0, T1 and T2) from each individual, grown overnight in duplicate and stocked. In parallel, phage populations were prepared from the same faecal samples by homogenising 30% w/v of faecal sample in phosphate buffered saline which was then centrifuged for 30 mins at 4,000 rpm and filter sterilised first through a 0.45 μm filter and then a 0.22 μm membrane pore filters to remove any bacteria but retain any phages present in the sample supernatant. These faecal phage populations (FPP) were stored at 4 °C and used within 24 hours.

### Time-shift assays and measures of local adaptation of bacterial x phage population interactions

Time-shift analysis of bacteria and phage populations was performed for all subjects in this study, n = 14. To do so, we tested the infectivity of the T1 phage population filtrate (FPP) against each of the 32 isolates sampled from that same individual for each time-point e.g. past (T0), contemporary (T1) and future bacterial hosts (T2), and conversely the sensitivity/resistance of bacteria sampled from T1 was tested against past (T0) and future (T2) phage population filtrates (FPPs) obtained from the same individual. For each study participant bacterial colonies (32 bacteria from each time-point, n=96) were grown overnight in LB broth. 300ul of FPP from T1 was added to each of the well which contained an individual bacterial hosts and the contents of each well were then added (independently) to a separate Falcon tube containing 5 ml of cooled (50 °C) molten soft agar (0.5% w/v) supplemented with Ca2+ and Mg2+ ions, mixed by swirling and then overlaid on a LB agar plate and incubated overnight at 37 °C. This analysis was conducted for all bacterial hosts from each study participant resulting in a total of 2240 ((96*14) + (32*2*14)) potential bacterium x phage population interactions. To test whether phage populations or bacteria were locally adapted *i.e.* more infective or resistant to their sympatric bacteria and phages respectively (i.e. sympatry or home = isolates from the same subject at the same time-point) compared to allopatric bacteria or phages that were isolated from another subject (away), we performed cross-inoculation assays using the home versus away measure of local adaption. For this analysis we assayed within-age group (infant v infant or adult v adult), and individuals were randomly paired. We only chose one sample-time-point per individual and for each pair of study subjects we tested phages taken from one subject against their own 32 bacterial hosts (Home) and 32 bacterial hosts from their paired subject (Away) and vice versa resulting in a “Home” and “Away” assay for each individual using soft agar overlay assays as outlined earlier. Of note T0 and T0 or T2 versus T2 combinations were used to extend the possible number of additional novel phage population x genotype interactions analysed in this study resulting in an additional 896 bacterium x phage population interactions. In all assays phage infectivity was scored as positive if we observed individual phage plaques (that are characteristic of phage infection, see fig. S8) against a given bacterial host. Phage plaques against sympatric hosts were isolated and stocked according to standard procedure. In parallel with these assays each FPP (n = 42) was also assayed against two *E. coli* indicator hosts namely *E. coli* MAC and *E. coli C* that are widely used to screen for the presence coliphages (*4*, *22*) using the same methodology for time-shift and LA assays described above.

### Individual phage infectivity x sympatric hosts for individuals I2 and A4

To disambiguate between population level and individual phage genotype infectivity against sympatric naturally occurring hosts over time we performed additional individual phage isolate x single bacterial host time-shift analysis for the cases of phage infectivity observed for individual I2 and individual A4 in our phage population time-shift and LA screens. Here four phages randomly selected and isolated from different sympatric bacterial hosts sampled at T1 from I2 and two phages isolated from two different sympatric bacterial hosts at T0 for A4 were amplified and tested against all sympatric temporally sampled bacterial hosts (n = 96) from I2 and A4, respectively. This analysis was performed using soft agar assays by spotting 10ul of each phage onto each of the 96 hosts sampled from either individual I2 or A4. All four phages isolated from individual I2 and both phages isolated from individual A4 had the same infectivity profile, respectively, across all hosts indicating that they were phenotypically identical with respect to host range.

### *In vitro* co-culture modelling using sympatric bacteria-temperate phages pairs isolated from individuals I2 and A4

The isolation of sympatric bacteria-phage pairs from I2 and A4 was exploited to investigate the temporal dynamics of bacteria and phage interactions and the genetic basis of resistance evolution to temperate phages. Sequence analysis of our chosen phages ph_I2_188 and phage ph_A4_364, isolated from individual I2 and A4 respectively, revealed that both were novel temperate phages (fig. S3, table S3), however, when both were BLASTED against the NCBI database both align partially to different phage contigs from a human metagenome study of 1000s of phages (*50*). These phages, together with a sequenced representative of the phage sensitive bacterial ST isolated from individual I2, designated I2_T1_19, and a sequenced representative of the phage sensitive *E. cloacae* strain isolated from individual A4, designated A4_T0_8, were used to establish our *in vitro* co-culture models. For each phage x bacteria combination, we set up in four replicate populations with 10^8^ bacteria and 10^6^ phages used to establish each population in 15ml Falcon tubes containing 6 ml of LB broth. Populations were incubated statically in an anaerobic chamber at 37 °C. To mimic more natural population turnover conditions in the human gut we established a protocol whereby we removed 2/3 of the population every 24 hours and replaced the volume removed (4ml) with fresh LB. Both experiments were run for a total of ten transfers or ten days (∼70 bacterial generations). Bacterial population densities were enumerated by serial dilution and plating, and phage populations were enumerated at T5 and T10 by serial dilution and spot plating on soft agar overlay using the ancestral host that was used to initiate the experiment. To test for potential co-evolution, we used time-shift analysis where filtrates of phage populations from Transfer 5 were tested for infectivity against 16 randomly selected bacterial isolates from T5 (n=16) and T10 (n=16) as well as the ancestral host (n=1) for each population and treatment, and conversely all bacteria from T5 were tested for resistance to the ancestral phage as well as sympatric phage populations from T5 and T10. We also tested T10 bacteria (n = 16 from each population for each model) for resistance to sympatric T10, T5 and ancestral phage. Phage infectivity was tested by spot-plating 10µl of phage filtrate from the relevant time-point and population onto soft-agar overlays of each bacterial host as appropriate to the time-shift assay.

### Identification and analysis of SPI phenotype in evolved resistant mutants

Lysogenised mutants of *E. coli* I2_T1_19 (n = 10) and *Enterobacter cloacae* A4_T0_8 (n=11) that evolved in our *in vitro* models were all evaluated for SPI phenotype. Here, single colonies were streaked to purity, grown overnight in LB broth and supernatants were filtered, serially diluted, and spotted onto soft agar overlays containing the ancestral host and hosts from whence they were derived. Plates were incubated overnight at 37 °C and filtered supernatants from overnight cultures of receptor mutants and the ancestral strains were also analysed as controls.

### Costs of resistance assays

We calculated ODmax to determine any potential bacterial fitness costs owing to phage resistance evolution. To do so, overnight cultures of I2_T1_19 and A4_T0_8 and their phage resistant mutants were serially diluted and 10ul of each dilution was inoculated into 190 µl of LB broth in 96-well plates in triplicate. Plates were incubated at 37 °C for 24 hours (H) and ODmax was calculated according to OD24H-OD0H values. Data for phage resistant mutants of A4_T0_8 and I2_T1_19 were analysed separately and were partitioned into lysogen or receptor modification as the basis of resistance and compared to WT growth over the same time-period using unpaired (between mutant class) and one-sample t-tests (mutant class versus WT) with a two-tailed distribution with statistical analysis carried out using IBM SPSS software.

### Bacterial strain and species identification within and between hosts

We used a combination of strain-typing PCRs, 16S rRNA Sanger sequencing and whole genome sequence analysis to capture the genetic diversity of and differentiate within and between *E. coli* and non-*E. coli Enterobacterales* species and strains isolated from individuals in this study. For all 96 isolates from each individual in this study (14*96 = 1344 isolates) we performed colony strain-genotyping PCRs. To do so, freshly grown overnight cultures of stocked strains were diluted 1/10 v/v in 200ul of 10% Chelex in a 96 well PCR plates and heated to 95 °C for 10 minutes. For strain-level fingerprinting 2µl of each diluted culture was added to a PCR master mix containing the strain-genotyping primer (CGG)4 (*51*) to discriminate between individual strains in a sample and assess strain level variation within an individual over the three timepoints sampled. PCR conditions were as per described elsewhere (*51*). PCR products were run on a 1.2% agarose gel for 90 minutes at 100V. Data from PCR strain profiling was used to pick representatives of unique strain profiles identified within each individual over time for 16S rRNA colony PCR and Sanger sequencing analysis followed by whole genome sequencing for a subset of strains from each individual as well as sensitive and resistant bacterial hosts from I2 and A4 that were identified in individual phage infectivity assays.

### SPI phenotype screening of bacterial isolates from the human gut and analysis of sympatric host resistance

We screened 96 bacterial hosts isolated from each of the adult and infant human hosts (n = 7) where phages were detected in at least one of the three time-points for an SPI+ phenotype using indicator *E. coli* hosts MAC and C (n = 672*2). For any unique SPI phenotype identified in this assay at least one representative of that SPI strain was grown overnight, the supernatant was filtered and 10ul of filtrate was then tested against all 96 sympatric bacteria from that same individual using soft agar overlays as previously described. As a control we also screened bacteria for an SPI from faecal samples where no phages were detected on any bacterial hosts including indicator hosts. To do so we sub-sampled the 96 hosts from each of these individuals based on strain typing profiles and tested 49 individual bacterial isolates representative of diversity present in samples analysed. For each bacterial host, each culture was grown overnight and 10 µl of filter sterilized supernatant was spotted onto soft agar overlays of our indicator hosts *E. coli* MAC and C as described earlier.

### Phage infectivity and bacterial resistance range analysis

To test for allopatric infectivity and phage host range together with the resistance range of bacteria isolated from the human gut we tested phages (n = 6) produced from naturally occurring SPI bacteria that were isolated and purified on indicator host MAC, together with phage A4_364 and phage I2_188 against 38 bacterial strains that we sub-sampled from our collection of isolates from the fourteen human hosts in this study. Strains were selected based on the PCR strain-typing analysis followed by whole genome sequencing such that the collection broadly represented the diversity of bacterial strains isolated in this study. In addition to the phages that were isolated from SPI hosts we also included a panel of model lytic phages T4, T5, T6 T7 and PP01 to provide a broader insight and comparative analysis of the resistance profiles of bacteria in the human gut. These model phages are relatively well characterised, represent different phage families and have been used in many experimental models of bacteria-phage interactions including studies of bacteria-phage coevolution. In the first set of experiments phages numbers were normalised to 10^5^ PFUs/ml to maintain biological realism as this was calculated as representative of the average and upper range of phages produced by SPI phenotypes isolated from faecal samples in our study (n = 6) were tested for phages produced per ml by growing overnight, serially diluting the supernatant and testing against the indicator host MAC-the average PFUs/ml formed on MAC was 1.28 x 10^5^; range 1.04 x 10^3^ – 2.85 x 10^5^. Phage host range/bacterial resistance assays were conducted using soft agar overlays where 10ul of phages were spotted onto each bacteria host in triplicate in two separate experiments. We followed up this experiment with a second assay, using the same soft agar overlay approach, but spotted each phage in triplicate over a range of titres ranging between 10^6^-10^9^ PFUs/ml.

### DNA sequencing and comparative genomics of ancestral strains, resistant mutants and natural strain isolates

Genomic DNA from all isolates in this study was extracted using the GenElute™ Bacterial Genomic DNA Kit as per manufacturer’s instructions and sent for next-generation sequencing to SeqCenter (https://www.seqcenter.com/). Ancestral strains A4_T0_8 and I2_T1_19 were sequenced using both short-(Illumina NovaSeq 6000) and long-read (Oxford Nanopore) technologies, while hybrid reference assemblies were generated using Flye (*52*) by SeqCenter. All bacterial mutants and naturally occurring isolates were sequenced using Illumina NovaSeq 6000 for paired short reads of 2 x 151bp. All raw reads were made available on Box (https://www.box.com) by SeqCenter and downloaded to a local Linux server for processing and analysis. WGS read quality was assessed using FastQC (v0.11.5) and filtered and trimmed using Trimmomatic (v0.39)(*53*) with the following parameters: “SLIDINGWINDOW:4:20 MINLEN:50 LEADING:3 TRAILING:3 HEADCROP:10”. Draft genomes were assembled using SPAdes (v3.11.1)(*54*) and contigs < 500bp were excluded from assemblies. Contamination and completeness of each assembly was assessed using CheckM (v1.2.2)(*55*) where all draft genomes passed the thresholds for high-quality assemblies: contamination < 1% and completeness > 99%. Genome-genome similarity for each pair of genomes was calculated using two methods: FastANI (v1.34)(*56*) for core nucleotide identity and an in-house script based on protocols used by the GGDC (v2.1)(*57*) webtool. Orthofinder (v2.5.5)(*58*) was used to determine homologous gene presence or absence across strains. Genes were predicted and assigned functions using prokka (v1.13)(*59*) for the general annotation of each genome. More specific functional annotations for each genome were predicted and assigned using the following command line software: EZClermont (v0.7.0)(*60*) for *E. coli* phylotyping, PADLOC (v2.0.0)(*61*) for anti-phage defense systems, SerotypeFinder (v2.0.2) for O- and H-antigen types, and MLST (v2.23.0) for sequence types. The PHASTEST webtool(*62*) was used to predict both intact and incomplete prophage regions within genomes. SNPs, indels and HGT events conferring resistance to all mutants were identified using breseq (v0.36.0)(*63*) with default parameters. False positive SNP and short indel predictions were excluded by searching raw reads for each SNP/indel plus 10bp upstream and downstream, since a 21bp read subsequence is short enough to have sufficient read coverage if present and long enough not to occur by chance if it is in fact absent from the region homologous to that of the reference genome. HGT events such as intra-genomic IS insertion and homologous recombination were confirmed by analysing and parsing BLAST (v2.6.0+)(*64*) and Clustal Omega (v1.2.4)(*65*) output, region specific PCRs and Sanger sequencing. All eight phages involved in this study were annotated using the Galaxy webtool Pharokka(*66*) (for the rapid annotation of bacteriophage genomes) and genome annotation was visualized linearly using the online webtool, Clinker (https://cagecat.bioinformatics.nl/tools/clinker). Integrated prophages, ph_I2_188 and ph_A4_364, were visualized with flanking host genes since they were identified by comparing respective ancestral reference genomes to lysogenized mutants. The remaining six phages were sequenced and assembled by SeqCenter from isolated virions and, hence, were visualized by reordering and reorienting all genes relative to the terminase large subunit. All heatmaps were generated using the R package pheatmap (v1.0.12) in R (v4.2.1).

## Data availability

All sequence data have been deposited in the NCBI Sequence Read Archive (SRA) under the BioProject, PRJNA1309996. Genbank files for the phages isolated in this study are available on Figshare using the following link https://figshare.com/s/19421f81ee1cb1f706ac <u>Subhead</u>

## Acknowledgements

Thank you to Prof. Marie Agnes Petit for providing indicator E. coli strains MAC and C and thank you to all participants who provided samples and made this study possible.

## Author contributions

Conceptualization: PDS

Methodology: PDS

Investigation: NL, HBH, NA, CH, AH, PDS

Visualization: HBH, AH, PDS

Funding acquisition: PDS

Project administration: PDS

Supervision: PDS

Writing – original draft: PDS

Writing – review & editing: PDS, CH, AH

## Data and materials availability

All data are available in the main text or the supplementary materials, with the exception of sequence data which has been deposited in and available at NCBI Sequence Read Archive (SRA) under the BioProject, PRJNA1309996, and Genbank files for the phages isolated in this study available on Figshare using the following link https://figshare.com/s/19421f81ee1cb1f706ac

## Supplementary Materials

Materials and Methods

Figs. S1 to S8

Tables S1 to S8

References (50–66)

