## Supplementary Figures and Tables for "Bacterial resistance and co-existence with temperate bacteriophages in the human gut"

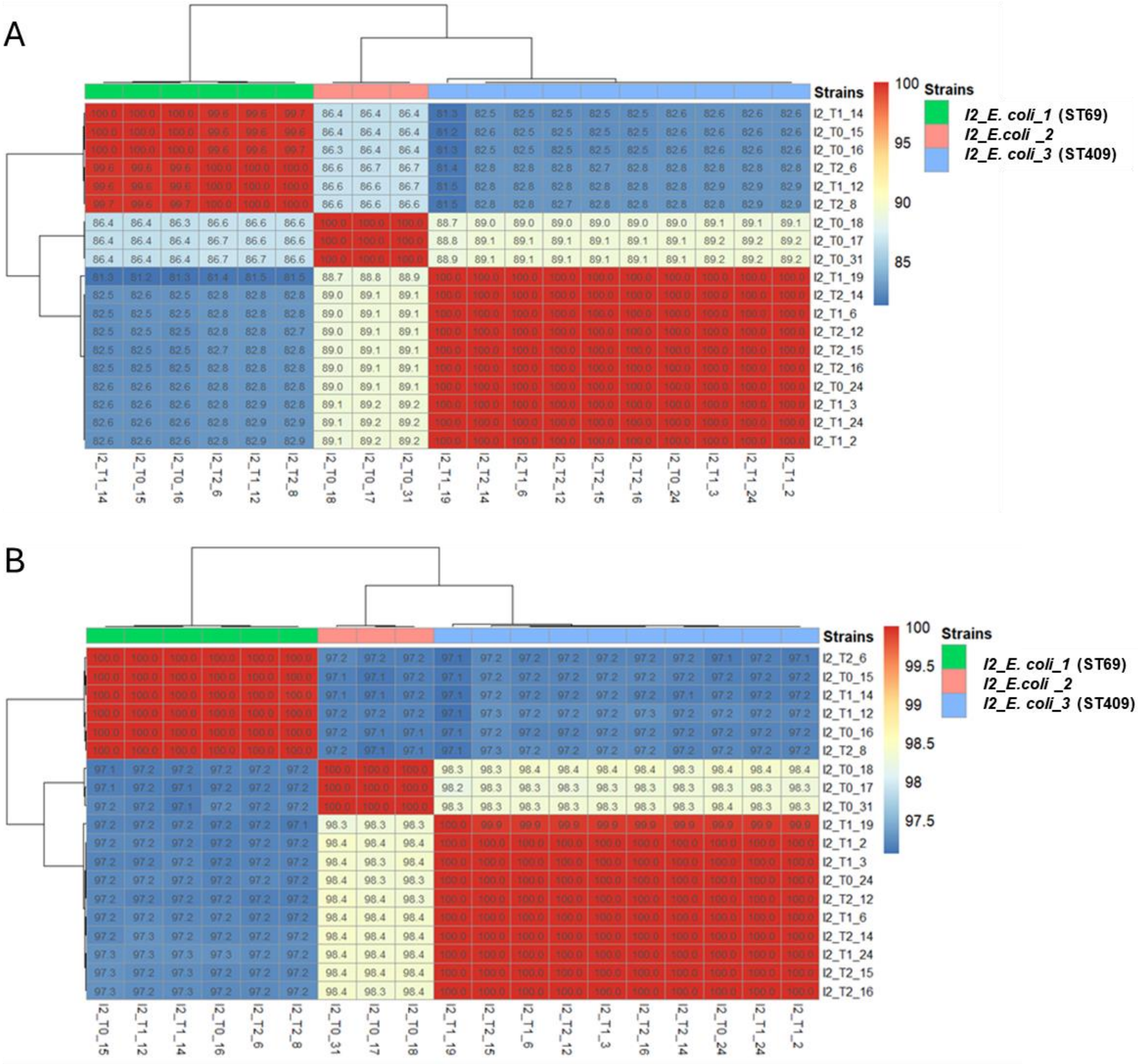

**Fig. S1. Heatmaps of representative isolates sampled from infant I2 clustered according to two common genome-genome similarity metrics. (A)** Average Nucleotide Identity (ANI) which measures sequence divergence in the core genome due to SNP level variation. **(B)** Genome Blast Distance Phylogeny (GBDP) which measures shared sequence homology as a percentage of total genome size.

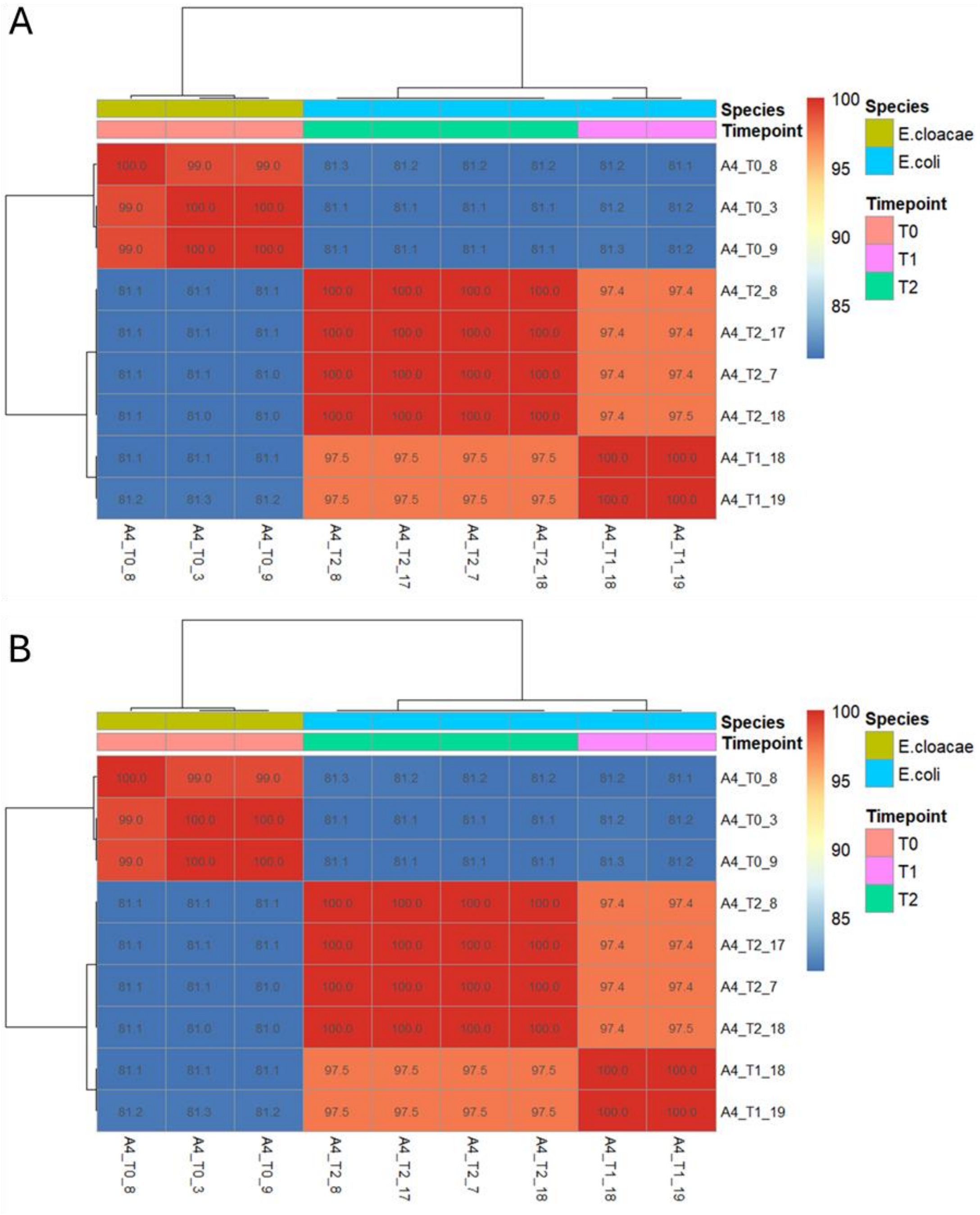

**Fig. S2. Heatmaps of representative isolates sampled from adult A4 clustered according to two common genome-genome similarity metrics.** (A) Average Nucleotide Identity (ANI) which measures sequence divergence in the core genome due to SNP level variation. (B) Genome Blast Distance Phylogeny (GBDP) which measures shared sequence homology as a percentage of total genome size.

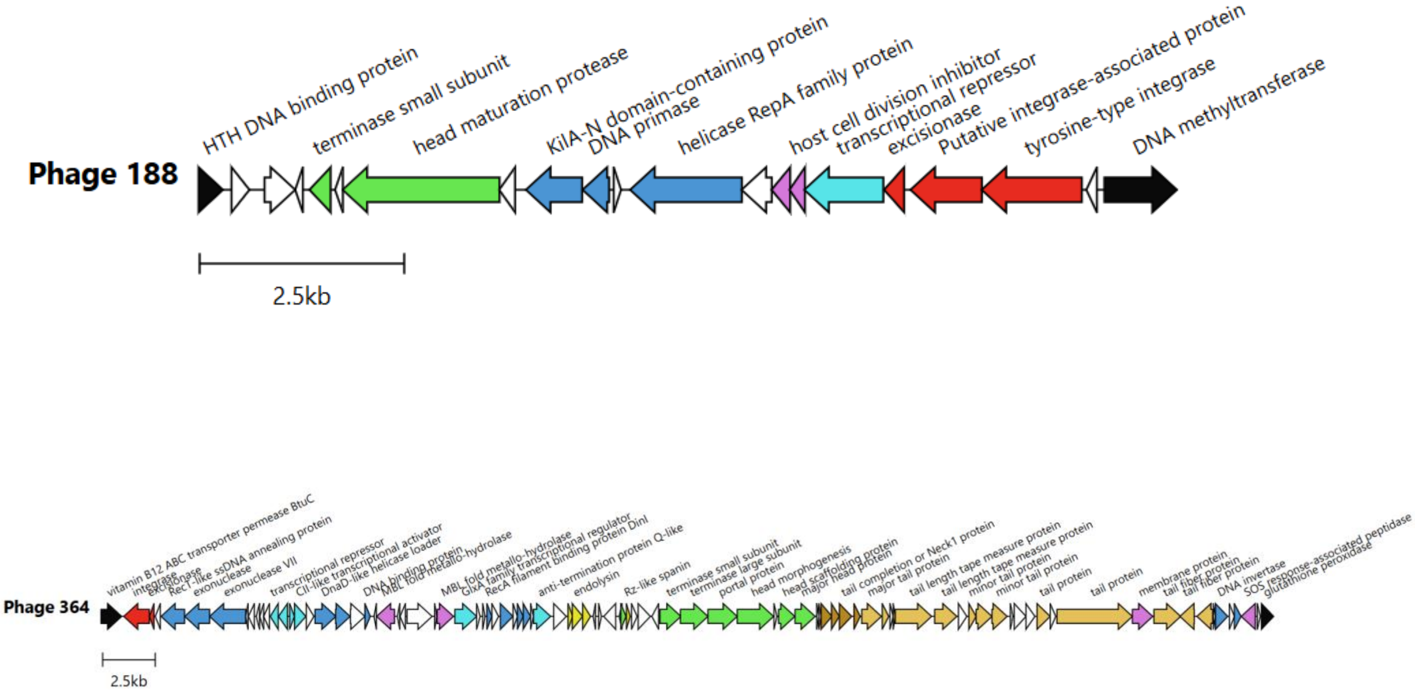

**Fig. S3. Genome organisation of ph_I2_188 and ph A4_364** (A) Phage I2_188 genome with integration site in host genome I2_T1_19. (B)Phage A_364 with integration site in host genome A4_T0_8. Flanking genes of the host (coloured in black). Phage-specific functional gene groups are coloured according to the following scheme: Head and packaging (green), DNA (blue), transcriptional regulation (light blue), integration and excision (red), tail (orange), lysis (yellow), head-tail connector (brown), hypothetical (grey) and other (purple). Genes were annotated using PHAROKKA and visualised using Clinker.

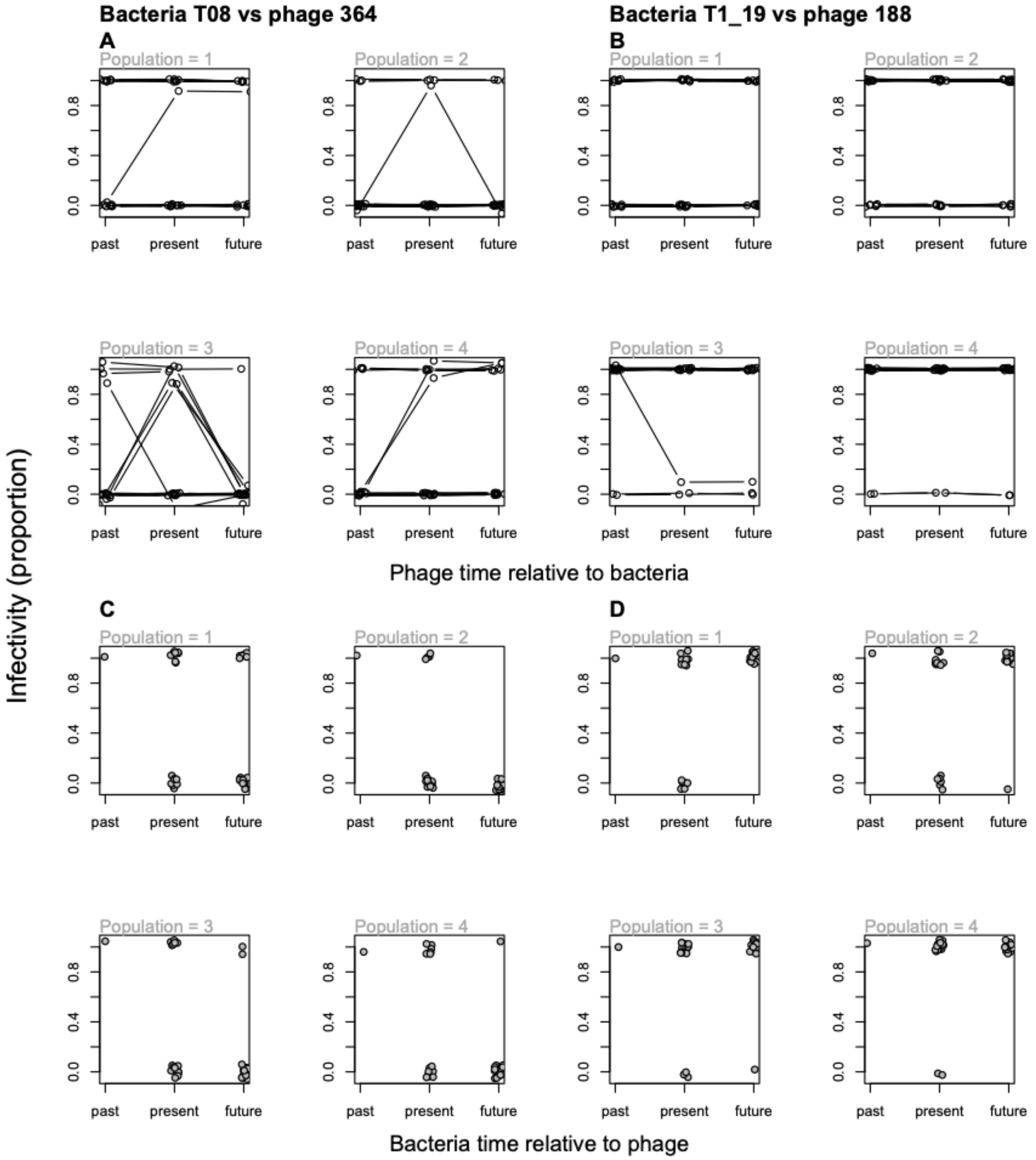

**Fig. S4. Infectivity profiles of bacterial colony isolates corresponding to population-level data shown in Fig. 2.** For each of the four panels of Fig. 2 (C-F), four subpanels are shown here (within each of A-D), one for each replicate population. S4 A and B: Infectivity profiles of 16 bacterial colony isolates (lines within each panel) from each of four populations in each of the two bacteria-phage combinations tested (labelled above each half of the figure). Each colony isolate was sampled from transfer 5, then tested for infectivity by phage lysates from the past (ancestral), present (transfer 5), or future (transfer 10). S4 C and D: Infectivity of phage lysates from transfer 5 in each of four populations for both bacteria-phage combinations. Points within panels show infectivity of phage lysate from transfer 5 against bacteria from the past (ancestral clone), middle (transfer 5; 16 colony isolates) or end (transfer 10; 16 colony isolates).

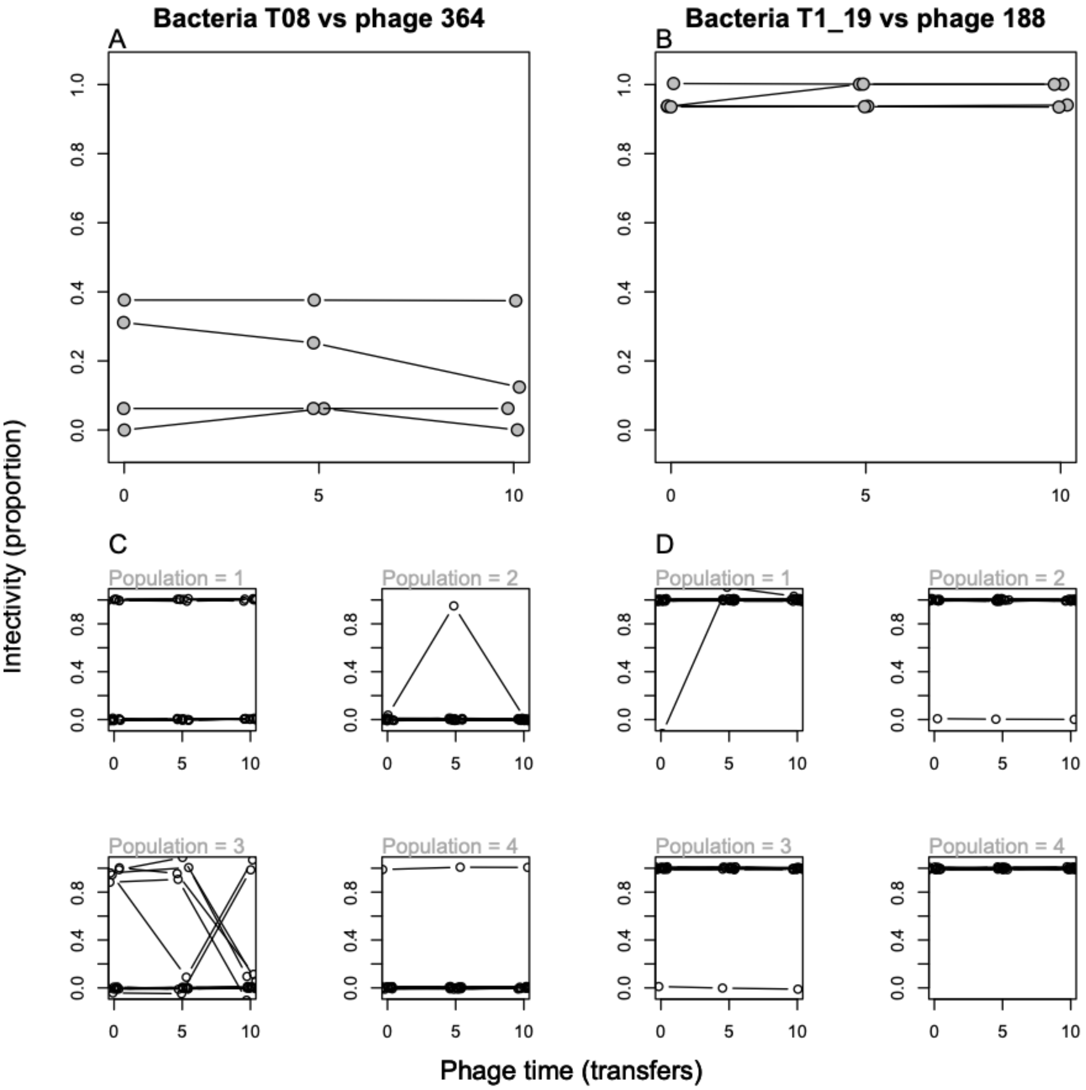

**Fig. S5.** A and B: Average infectivity (proportion of 16 bacterial colony isolates isolated at transfer 10 that could be infected) for phage lysates from the start (transfer 0), middle (transfer 5) and end (transfer 10) of the experiment. Lines within panels show four replicate populations in each bacteria-phage combination (labelled at top). As for bacteria sampled from transfer 5, the average slope of infectivity over phage time was not significantly different from zero in any population ($p$>0.05 when tested by one-sample $t$-tests with the 16 slope estimates, each from one bacterial colony isolate, done separately for each of the four replicate populations in each phage-bacterium combination). S5 C and D: Infectivity profiles of colony isolates used to calculate the population-level means shown in panels A and B. Infectivity profiles of 16 bacterial colony isolates (lines within each panel) from each of four populations (the four subpanels in each of C and D) are shown in each bacteria-phage combination. Each colony isolate was sampled from transfer 10, then tested for infectivity by phage lysates from the start (ancestral), middle (transfer 5), or end (transfer 10) of the experiment.

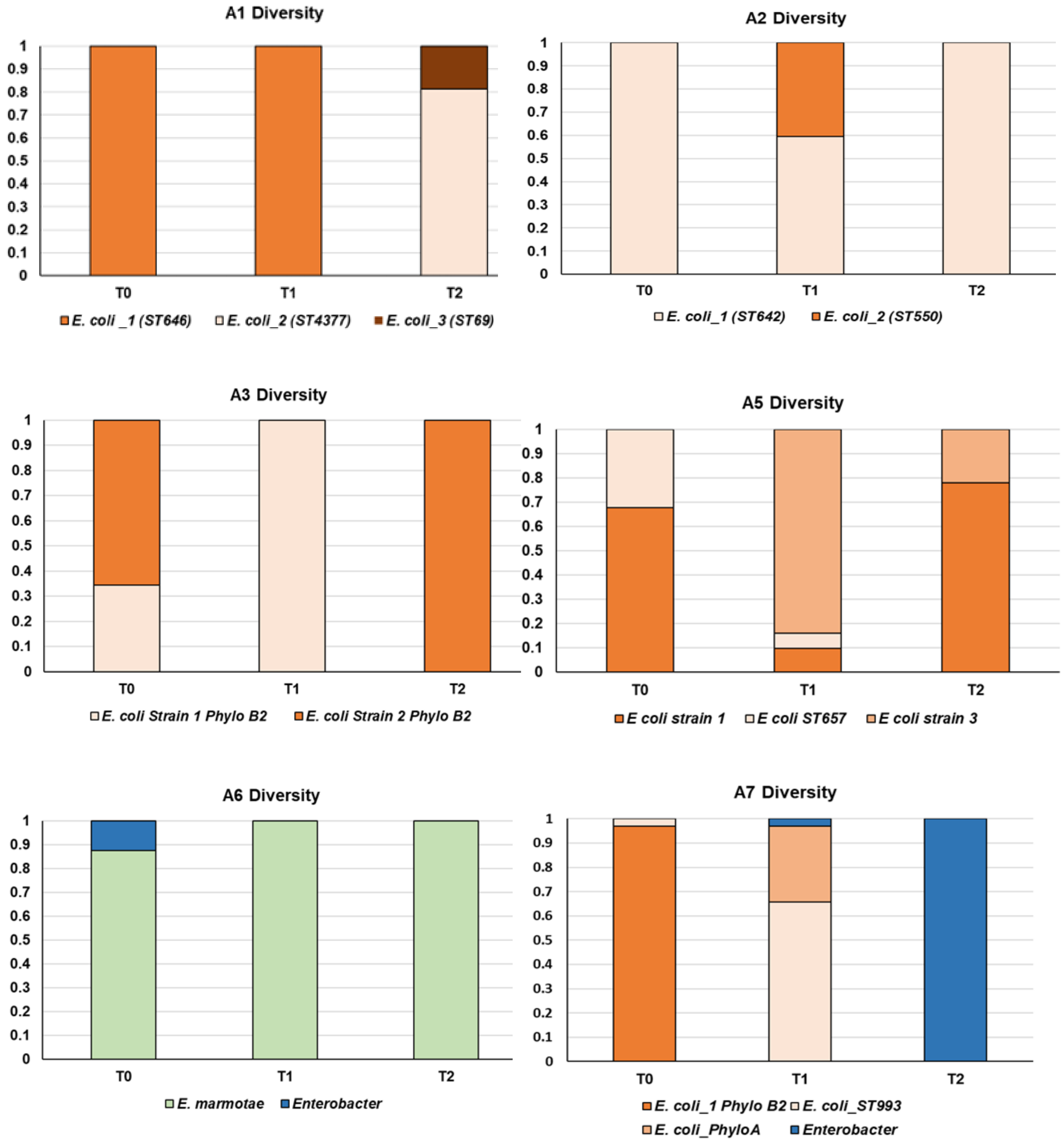
A1 Diversity
T0
T1
T2
E. coli _1 (ST646)
E. coli_2 (ST4377)
E. coli_3 (ST69)
A2 Diversity
T0
T1
T2
E. coli_1 (ST642)
E. coli_2 (ST550)
A3 Diversity
T0
T1
T2
E. coli Strain 1 Phylo B2
E. coli Strain 2 Phylo B2
A5 Diversity
T0
T1
T2
E coli strain 1
E coli ST657
E coli strain 3
A6 Diversity
T0
T1
T2
E. marmotae
Enterobacter
A7 Diversity
T0
T1
T2
E. coli_1 Phylo B2
E. coli_ST993
E. coli_PhyloA
Enterobacter

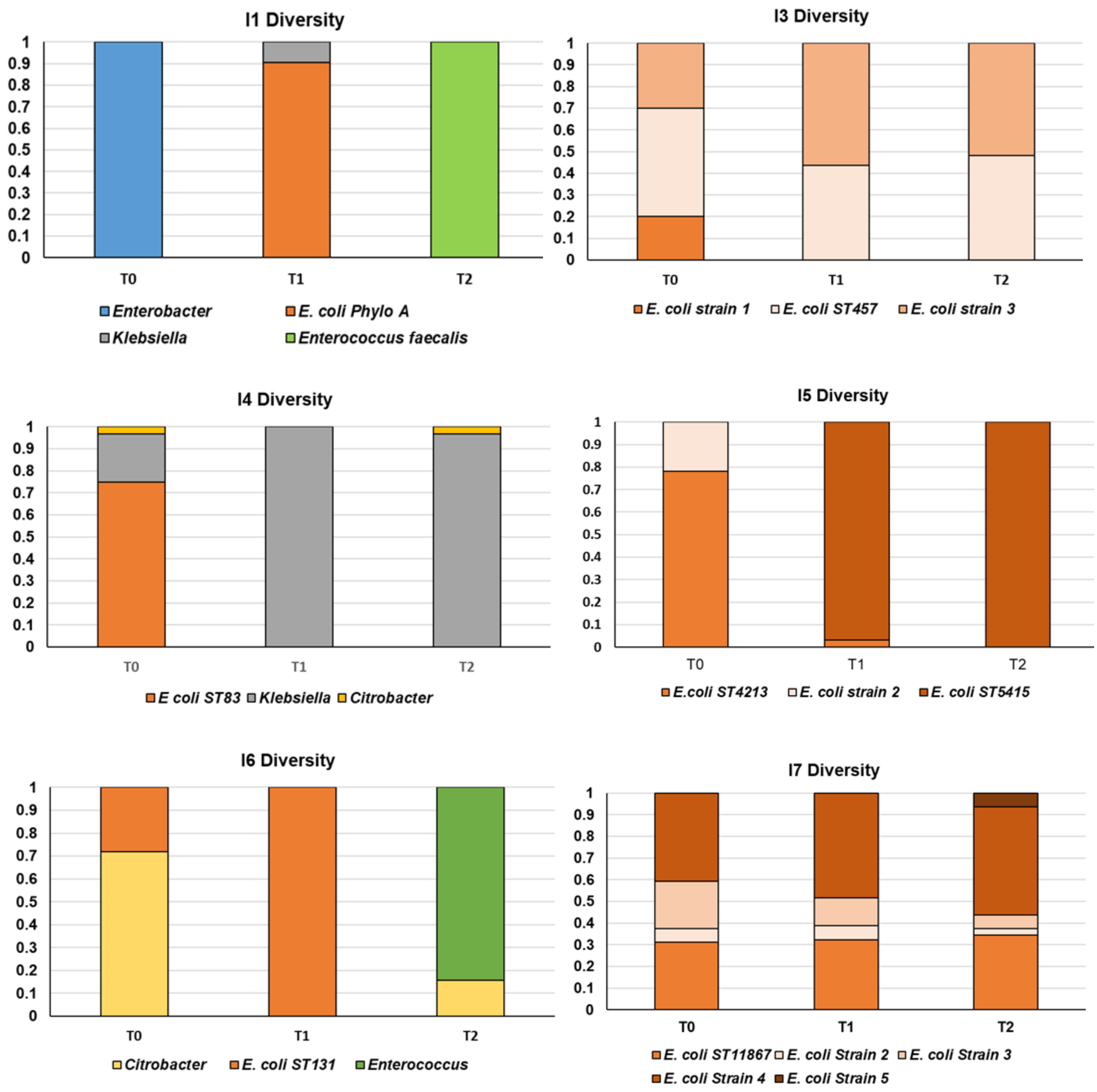

**Fig. S6. Temporal analysis and relative abundance of bacterial diversity for each individual in this study showing genus, species and strain level variation within individuals over time and between individual hosts.** Note taxonomic identification of bacteria was obtained from both 16S rRNA PCR and sequencing of unique strain types identified *via* strain typing PCRs and whole genome sequencing in many cases. Where *E. coli* ST designations are given, this is based on WGS data.

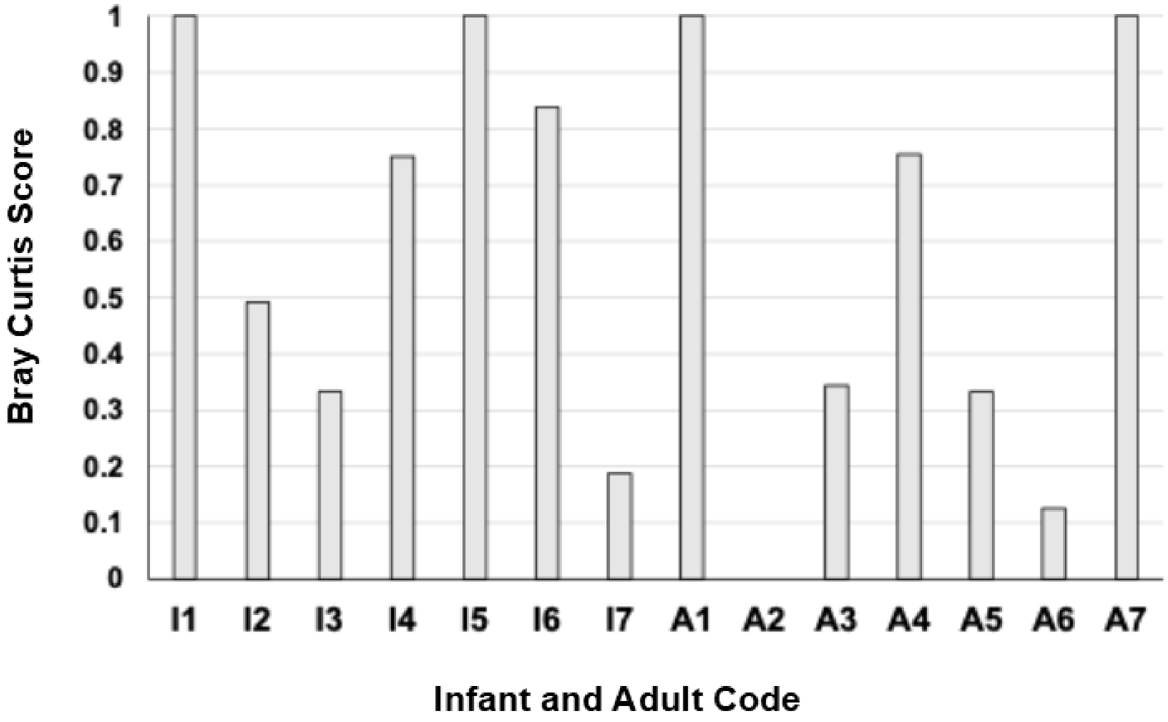

**Fig. S7. Bray Curtis dissimilarity analysis of *Enterobacterales* diversity within individuals between time-points T0 and T2.** Scale ranges from 0 to 1 indicating proportion of dissimilarity between these two timepoints with a score of 0 = identical similarity and 1 = no similarity.

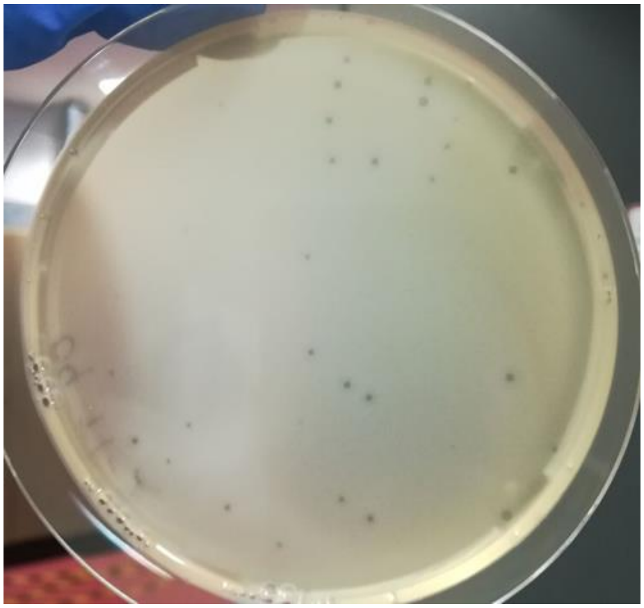

**Fig S8**. **Example of a positive case of phage infectivity against a sympatric host in faecal phage filtrate phage population analysis.** Evidence for phages infecting host I2_T1_19 in our FPP phage population analysis screen with individual plaques on this host showing positive phage infectivity in sympatry for this phage population x individual host interaction.

**Supplementary Table 1.** Taxonomic analysis and affiliations of phages isolated and sequenced in this study

| Phage | Realm | Kingdom | Phylum | Class | Order | Family | Subfamily | Genus | Species | Top NCBI hit | % query coverage | % identity |
|---|---|---|---|---|---|---|---|---|---|---|---|---|
| I2_ph_188 | Unknown | Unknown | Unknown | Unknown | Unknown | Unknown | Unknown | New_genus | New_species | Caudoviricetes sp. isolat | 40 | 97 |
| A4_ph_364 | Unknown | Unknown | Unknown | Unknown | Unknown | Unknown | Unknown | New_genus | New_species | Caudoviricetes sp. isolat | 23 | 92.9 |
| SPI_A5_02 | Duplodnaviria | Heunggongvirae | Uroviricota | Caudoviricetes | Not Defined Yet | Not Defined Yet | Not Defined Yet | Nesevirus | Nesevirus new_name | | | |
| SPI_A7_57 | Unknown | Unknown | Unknown | Unknown | Unknown | Unknown | Unknown | New_genus | New_species | Caudoviricetes sp. isolat | 28 | 94.9 |
| SPI_I2_4 | Duplodnaviria | Heunggongvirae | Uroviricota | Caudoviricetes | Not Defined Yet | Not Defined Yet | Not Defined Yet | Jouyvirus | Jouyvirus new_name | | | |
| SPI_A1_69 | Duplodnaviria | Heunggongvirae | Uroviricota | Caudoviricetes | Not Defined Yet | Not Defined Yet | Not Defined Yet | Jouyvirus | Jouyvirus jv1H12 | | | |
| SPI_A1_62 | Duplodnaviria | Heunggongvirae | Uroviricota | Caudoviricetes | Not Defined Yet | Peduoviridae | Not Defined Yet | Peduovirus | Peduovirus new_name | | | |
| SPI_A2_90 | Duplodnaviria | Heunggongvirae | Uroviricota | Caudoviricetes | Not Defined Yet | Peduoviridae | Not Defined Yet | Peduovirus | Peduovirus new_name | | | |

**Supplementary Table 2.** Summary of mutational events, and loci and genes affected for evolved bacerial resistant mutants in vitro co-culture experiments with A4 T0_8 and I2_T1_19.

| **A4_T0_08** | **Resistance mechanism** | **Prophage status** | **Prophage or IS integration site** | **Gene name** | **Gene function** | **Mutation type** | **Mutation locus** |
|---|---|---|---|---|---|---|---|
| **1_1_R** | Lysogenic conversion | Integrated | Vitamin B12 import system permease protein BtuC/Glutathione peroxidase BtuE | MKEMDIJK_01759/MKEMDIJK_01760 | Vitamin B12 import | Integration/site-specific recombination | 3'-end of gene: AAGCTGGACGTTAA -> AGGCTAATCGCTGA |
| **1_2_R** | Lysogenic conversion | Integrated | Vitamin B12 import system permease protein BtuC/Glutathione peroxidase BtuE | MKEMDIJK_01759/MKEMDIJK_01760 | Vitamin B12 import | Integration/site-specific recombination | 3'-end of gene: AAGCTGGACGTTAA -> AGGCTAATCGCTGA |
| **1_6_R** | Receptor modification | | | MKEMDIJK_02705 | Glycosyltransferase | Deletion: A | residue 378 : AAAATGA -> AAATGA |
| **2_1_R** | Lysogenic conversion | Integrated | Vitamin B12 import system permease protein BtuC/Glutathione peroxidase BtuE | MKEMDIJK_01759/MKEMDIJK_01760 | Vitamin B12 import | Integration/site-specific recombination | 3'-end of gene: AAGCTGGACGTTAA -> AGGCTAATCGCTGA |
| **2_5_R** | Lysogenic conversion | Integrated | Vitamin B12 import system permease protein BtuC/Glutathione peroxidase BtuE | MKEMDIJK_01759/MKEMDIJK_01760 | Vitamin B12 import | Integration/site-specific recombination | 3'-end of gene: AAGCTGGACGTTAA -> AGGCTAATCGCTGA |
| **2_16_R** | Lysogenic conversion | Integrated | Vitamin B12 import system permease protein BtuC/Glutathione peroxidase BtuE | MKEMDIJK_01759/MKEMDIJK_01760 | Vitamin B12 import | Integration/site-specific recombination | 3'-end of gene: AAGCTGGACGTTAA -> AGGCTAATCGCTGA |
| **3_1_R** | Lysogenic conversion | Integrated | Vitamin B12 import system permease protein BtuC/Glutathione | MKEMDIJK_01759/MKEMDIJK_01760 | Vitamin B12 import | Integration/site-specific recombination | 3'-end of gene: AAGCTGGACGTTAA -> AGGCTAATCGCTGA |

| | | | | | | | |
|---|---|---|---|---|---|---|---|
| | | | peroxidase BtuE | | | | |
| **3_2_R** | Receptor modification | | | MKEMDIJK_02707 | O-antigen ligase | Insertion: A | residue 513: AATAGT -> AAATAG |
| **3_6_R** | Receptor modification | | | MKEMDIJK_02707 | O-antigen ligase | Insertion: 29 bp duplication | residue 756: TGCACTAAGTGGTTATTTCATAAATAAAATGCA |
| **4_12_R** | Receptor modification | | | MKEMDIJK_02707 | O-antigen ligase | Deletion: A | residue 350: AAAAAAA -> AAAAAA leads to TGA at residue 367 |

| **I2_T1_19** | **Resistance mechanism** | **Prophage status** | **Prophage or IS integration site** | **Gene name** | **Gene function** | **Mutation type** | **Mutation locus** |
|---|---|---|---|---|---|---|---|
| **R.1.6** | Lysogenic conversion | Integrated | DNA adenine methyltransferase /DNA-binding protein Fis | OJJPFJGE_00488/OJJPFJGE_00489 | DNA-binding | Integration/site-specific recombination | 3'-end of gene: AAAATACGGCATGAACTAA -> AAAATACGGCATGAACTGA |
| **R.1.10** | Lysogenic conversion | Integrated | DNA adenine methyltransferase /DNA-binding protein Fis | OJJPFJGE_00488/OJJPFJGE_00489 | DNA-binding | Integration/site-specific recombination | 3'-end of gene: AAAATACGGCATGAACTAA -> AAAATACGGCATGAACTGA |
| **R.1.15** | Receptor modification | | | OJJPFJGE_01758-OJJPFJGE_01773 | PMM operon | Large deletion due to homologous recombination | OJJPFJGE_01758-OJJPFJGE_01773 |
| **R.2.1** | Receptor modification | | | OJJPFJGE_01758-OJJPFJGE_01773 | PMM operon | Large deletion due to homologous recombination | OJJPFJGE_01758-OJJPFJGE_01773 |
| **R.2.11** | Lysogenic conversion | Circular contig | No integration (phagemid) | | | | |
| **R.2.12** | Lysogenic conversion | Circular contig | No integration (phagemid) | | | | |

| | | | | | | | |
|---|---|---|---|---|---|---|---|
| **R.2.15** | Lysogenic conversion | Integrated | DNA adenine methyltransferase /DNA-binding protein Fis | OJJPFJGE_00488/OJJPFJGE_00489 | DNA-binding | Integration/site-specific recombination | 3'-end of gene: AAAATACGGCATGAACTAA -> AAAATACGGCATGAACTGA |
| **R.3.11** | Receptor modification | | Glycosyl transferase (one gene upstream from O-antigen ligase) | OJJPFJGE_00080 | O-antigen biosynthesis | Integration/transposition of IS5 transposase | residue 297 of the gene OJJPFJGE_00080 |
| **R.3.15** | Receptor modification | | O-antigen ligase | OJJPFJGE_00081 | O-antigen biosynthesis | Integration/transposition of IS5 transposase | residue 934 of the gene OJJPFJGE_00081 |
| **R.3.4** | Receptor modification | | O-antigen ligase | OJJPFJGE_00081 | O-antigen biosynthesis | Integration/transposition of IS1 InsB | residue 1129 of the gene OJJPFJGE_00081 |

**Supplementary Table 3**. **Read mapping for PMM operon for I2_T1_19 ancestor and evolved resistant mutants.** Note for two mutants R1_1_15 and R_2_1 from two separate populations homologous recombination between two homologous genes resulted in the excision of 14 genes in the PMM operon, see also Figure 4 A and main text. Deletion was also confirmed by PCR analysis.

| Gene Locus | R_1_10 | R_1_15 | R_1_6 | R_2_11 | R_2_15 | R_2_1 | R_3_11 | R_3_15 | R_3_4 |
|---|---|---|---|---|---|---|---|---|---|
| **OJJPFJGE_01757** | 61.0883 | 62.6616 | 57.3085 | 57.2144 | 56.4333 | 127.176 | 119.896 | 62.981 | 59.38 |
| **OJJPFJGE_01758** | 64.2036 | 0 | 60.5849 | 66.2832 | 50.5971 | 0 | 138.119 | 57.848 | 57.053 |
| **OJJPFJGE_01759** | 66.0649 | 0 | 64.0974 | 59.4692 | 67.2299 | 0 | 125.463 | 55.192 | 63.3469 |
| **OJJPFJGE_01760** | 64.4239 | 0 | 67.5746 | 62.7572 | 59.1639 | 0 | 135.398 | 68.644 | 57.9688 |
| **OJJPFJGE_01761** | 61.6781 | 0 | 53.6806 | 56.6249 | 61.8788 | 0 | 125.279 | 58.7559 | 58.1638 |
| **OJJPFJGE_01762** | 68.8143 | 0 | 63.5857 | 55.2803 | 50.6753 | 0 | 124.465 | 70.7283 | 59.8774 |
| **OJJPFJGE_01763** | 57.2875 | 0 | 63.0694 | 53.3949 | 52.1398 | 0 | 104.442 | 51.8031 | 58.245 |
| **OJJPFJGE_01764** | 63.6436 | 0 | 59.988 | 52.8794 | 59.9715 | 0 | 111.867 | 57.0037 | 48.6621 |
| **OJJPFJGE_01765** | 72.2467 | 0 | 68.1533 | 62.2456 | 54.7844 | 0 | 124.856 | 55.5689 | 60.9356 |
| **OJJPFJGE_01766** | 57.4061 | 0 | 56.8635 | 55.7258 | 51.868 | 0 | 118.704 | 45.5859 | 57.5597 |
| **OJJPFJGE_01767** | 46.404 | 0 | 44.0489 | 51.125 | 50.4293 | 0 | 87.0453 | 46.1793 | 50.9601 |
| **OJJPFJGE_01768** | 43.7509 | 0 | 53.2708 | 53.2518 | 52.2509 | 0 | 112.075 | 57.5625 | 50.6033 |
| **OJJPFJGE_01769** | 50.8547 | 0 | 54.6024 | 60.0512 | 48.297 | 0 | 108.44 | 54.3501 | 48.6611 |
| **OJJPFJGE_01770** | 63.9062 | 0 | 47.3037 | 42.3889 | 40.8037 | 0 | 99.0099 | 53.3383 | 52.9148 |
| **OJJPFJGE_01771** | 51.677 | 0 | 46.6305 | 45.7959 | 43.1744 | 0 | 96.4328 | 49.832 | 54.1757 |
| **OJJPFJGE_01772** | 89.6996 | 29.4688 | 95.466 | 89.113 | 76.0835 | 57.8639 | 188.451 | 103.693 | 90.746 |
| **OJJPFJGE_01773** | 67.5062 | 5.10503 | 57.2458 | 62.1583 | 53.8023 | 7.28446 | 127.446 | 58.4143 | 61.0153 |

**Supplementary Table 4.** Genes presence/absence and similarity between I2_T1_19 and two representatives of co-occurring phage resistant *E. coli* strains isolated from individual I2, namely I2_T0_15 and I2_T0_17

| | **I2_T0_15** | | | | **I2_T0_17** | | | | |
|---|---|---|---|---|---|---|---|---|---|
| **I2_T1_19 genes** | **% identity** | **% query** | **SNPs** | **indels** | **% identity** | **% query** | **SNPs** | **indels** | **Functional annotation** |
| **OJJPFJGE_00073** | 97.67 | 97.73 | 8 | 0 | 99.13 | 97.73 | 3 | 0 | lipopolysaccharide core heptosyl transferase III |
| **OJJPFJGE_00074** | 98.93 | 100 | 4 | 0 | 99.2 | 100 | 3 | 0 | lipopolysaccharide glucosyltransferase I |
| **OJJPFJGE_00075** | 99.62 | 100 | 1 | 0 | 100 | 100 | 0 | 0 | lipopolysaccharide core heptose (I) kinase |
| **OJJPFJGE_00076** | 54.25 | 90.53 | 139 | 1 | 99.41 | 100 | 2 | 0 | UDP-D-glucose:(glucosyl)LPS alpha-1,3-glucosyltransferase |
| **OJJPFJGE_00077** | 43.69 | 93.35 | 174 | 0 | 99.7 | 100 | 1 | 0 | UDP-glucose:(glucosyl)LPS alpha-1,2-glucosyltransferase |
| **OJJPFJGE_00078** | 53.81 | 91.3 | 97 | 0 | 99.08 | 94.35 | 2 | 0 | lipopolysaccharide core heptose (II) kinase |
| **OJJPFJGE_00079** | 38.89 | 95.01 | 184 | 6 | 99.12 | 100 | 3 | 0 | UDP-glucose:(glucosyl)LPS alpha-1,2-glucosyltransferase |
| **OJJPFJGE_00080** | 33.33 | 65.14 | 132 | 6 | 97.86 | 100 | 7 | 0 | Undecaprenyl-phosphate 4-deoxy-4-formamido-L-arabinose transferase |
| **OJJPFJGE_00081** | 34.15 | 9.83 | 26 | 1 | 99.52 | 100 | 2 | 0 | O-antigen ligase |
| **OJJPFJGE_00082** | 95.19 | 96.59 | 15 | 0 | 99.07 | 100 | 3 | 0 | ADP-heptose:LPS heptosyltransferase I |
| **OJJPFJGE_00083** | 99.7 | 95.98 | 1 | 0 | 99.7 | 96.26 | 1 | 0 | ADP-heptose:LPS heptosyltransferase II |
| | Missing in I2_T0_15 | | | | | | | | |

**Supplementary Table 5.** Genes presence/absence and similarity between A4_T0_8 and one representative of co-occurring phage resistant *E. cloacae* strain isolated from individual A4, namely A4_T0_T3

| | **A4_T0_03** | | | | |
|---|---|---|---|---|---|
| **A4_T0_08** | **% identity** | **% query** | **SNPs** | **indels** | **Functional annotation** |
| MKEMDIJK_02700 | 97.68 | 100 | 9 | 0 | UDP-glucose 6-dehydrogenase |
| MKEMDIJK_02701 | 64.21 | 97.27 | 102 | 0 | Glucose-1-phosphate thymidylyltransferase |
| MKEMDIJK_02702 | 67.14 | 97.78 | 104 | 2 | dTDP-glucose 4,6-dehydratase |
| MKEMDIJK_02703 | 98.93 | 100 | 5 | 0 | 6-phosphogluconate dehydrogenase, decarboxylating |
| MKEMDIJK_02704 | 30.43 | 33.95 | 61 | 2 | UDP-Gal:alpha-D-GlcNAc-diphosphoundecaprenol beta-1,3-galactosyltransferase |
| MKEMDIJK_02705 | no hit | NA | NA | NA | beta-1,6-N-acetylglucosaminyltransferase |
| MKEMDIJK_02706 | 34.04 | 16.04 | 31 | 0 | glycosyltransferase family 2 protein |
| MKEMDIJK_02707 | 24.78 | 28.9 | 63 | 5 | O-antigen ligase |
| MKEMDIJK_02708 | 25.64 | 18.44 | 56 | 1 | oligosaccharide flippase |
| MKEMDIJK_02709 | 50 | 8.91 | 11 | 0 | glycosyltransferase |
| MKEMDIJK_02710 | 97.29 | 99.33 | 8 | 0 | UTP--glucose-1-phosphate uridylyltransferase |
| MKEMDIJK_02711 | 29.86 | 107.25 | 195 | 19 | N-acetyl-alpha-D-glucosaminyl-diphospho-ditrans,octacis-undecaprenol 4-epimerase |
| MKEMDIJK_02712 | 98.92 | 100 | 5 | 0 | colanic acid biosynthesis protein WcaM |
| | Missing in A4_T0_03 | | | | |

**Supplementary Table 6.** Phage infectivity* against *Enterobacterales* isolates representing diversity of genera and species present in dataset together with sequence type, and H and O-antigen information where appropriate

| Code | Phylogroup or Species I | ST | H | O antigen | SPI_A1_62 | SPI_A1_69 | SPI_A2_90 | SPI_A5_02 | SPI_A7_57 | SPI_I2_4 | I2_ph_188 | A4_ph_364 | T4 | T5 | T6 | T7 | PPO1 |
|---|---|---|---|---|---|---|---|---|---|---|---|---|---|---|---|---|---|
| **A1_T1_13** | **B2** | ST646 | fliC_H6 | wzx_O166 wzy_O166 | 0 | | 0 | 0 | 0 | 0.00025 | 0 | 0 | 0.0086 | | 0.067 | 0.07 | 0.0003 |
| **A1_T1_8** | **B2** | ST646 | fliC_H6 | wzx_O166 wzy_O166 | 0 | | 0 | 0 | 0 | | 0 | 0 | 0 | 0 | 0.053333 | 0 | 0 |
| **A1_T2_11** | **D** | ST4377 | fliC_H18 | NO_HITS_FOUND | 0 | 0 | 0 | 0 | 0 | 0 | 0 | 0 | 0 | 0 | 0 | 0 | 0 |
| **A1_T2_25** | **D** | ST69 | fliC_H18 | wzx_O17/O77 wzy_O17/O44 | 0 | 0 | | | | | | 0 | 0 | 0 | 0 | 0 | 0 |
| **A1_T2_4** | **D** | ST4377 | fliC_H18 | NO_HITS_FOUND | 0 | 0 | 0 | 0 | 0 | 0 | 0 | 0 | 0 | 0 | 0 | 0 | 0 |
| **A1_T2_5** | **D** | ST69 | fliC_H18 | wzx_O17/O77 wzy_O17/O44 | 0 | 0 | | | | | | 0 | 0 | 0 | 0 | 0 | 0 |
| **A2_T0_1** | **D** | ST69 | fliC_H1 | wzx_O15 wzy_O15 | 0 | 0 | 0 | 0 | 0 | 0 | 0 | 0 | 0 | 0 | 0 | 0 | 0 |
| **A2_T1_15** | **B1** | ST642 | fliC_H4 | wzx_O184 wzy_O184 | 0 | 0 | | | | 0 | | 0 | 0 | 0 | 0 | 0 | 0 |
| **A3_T1_15** | **B2** | ST550 | fliC_H5 | wzx_O75 wzy_O75 | 0 | 0 | 0 | 0 | 0 | 0 | 3 x 10^5 | 0 | 0 | 0 | 0.001063 | 0 | |
| **A3_T2_18** | **B2** | ST95 | fliC_H7 | wzx_O50 wzy_O2 | 0 | 0 | 0 | 0 | 0 | 0 | 0 | 0 | 0 | 0 | 0 | 0 | 0 |
| **A5_T0_24** | **G** | ST657 | fliC_H18 | wzx_O183 wzy_O183 | 0 | 0 | 0 | 0 | 0 | 0 | 0 | 0 | 0 | 0 | 0 | 0 | 0 |
| **A6_T0_4** | **A** | - | fliC_H48 | wzx_O16 wzy_O16 | 0 | 0 | 0 | 0 | 0 | 0 | 0 | 0 | 0 | 0 | 0 | 0 | 0 |
| **A6_T0_18** | ***E. marmotae*** | | fliC_H56 | wzx_O150 wzy_O150 | 0 | 7.51E-05 | 0.00354 | 0 | 0 | 0.0096 | 0 | 0 | 0 | | 0.008 | 0 | 0 |
| **A7_T0_11** | **A** | ST993 | fliC_H30 | wzx_O100 wzy_O100 | 0 | 0 | 0 | 0 | 0 | 0 | 0 | 0 | 0 | 0 | 0 | 0 | 0 |
| **A7_T1_7** | **A** | ST993 | fliC_H30 | wzx_O100 wzy_O100 | 0 | 0 | 0 | 0 | 0 | 0 | 0 | 0 | 0 | 0 | 0 | 0 | 0 |
| **A3 T0_4** | **B2** | - | fliC_H6 fliC | wzx_O166 wzx_O111 | 0 | 0 | 0 | 0 | 0 | | 0 | 0 | 0 | 0 | 0 | 0 | 0 |
| **I2_T1_19** | **A** | ST409 | fliC_H38 fl | wzx_O75 wzy_O75 | 0 | 0 | 0 | 0 | 0 | 0 | 4 x 10^6 | 0 | 0 | 0 | 0 | 0 | 0 |
| **I2_T1_25** | **D** | ST69 | fliC_H18 | wzx_O15 wzy_O15 | 0 | 0 | 0 | 0 | 0 | 0 | 0 | 0 | 0 | 0 | 0 | 0 | 0 |
| **I3_T1_1** | **F** | ST457 | fliC_H25 | wzx_O11 wzy_O11 | 0 | 0 | 0 | 0 | 0 | 0 | 0 | 0 | 0 | 0 | 0 | 0 | 0 |
| **I4_T1_25** | **B2** | ST83 | fliC_H5 | wzx_O25 wzy_O25 | 0 | 0 | 0 | 0 | 0 | 0.00012 | 0 | 0 | 2.86E-06 | | 0.000008 | 0 | |
| **I5_T0_12** | **B1** | | fliC_H21 | wzx_O55 wzy_O55 | 0 | 0 | 0 | 0 | 0 | | 0 | 0 | 0.0014 | 0.006 | 0.000013 | 0.067 | 0 |
| **I5_T2_29** | **B1** | ST5415 | fliC_H7 | wzx_O178 | 0 | 0 | 0 | 0 | 0 | 0 | 0 | 0 | 0 | 0 | 0 | 0 | 0 |
| **I6_T0_9** | **B2** | ST131 | fliC_H5 | wzx_O16 wzy_O16 | 0 | 0 | 0 | 0 | 0 | 0 | 0 | 0 | 0 | 0 | 0 | 0 | 0 |
| **I6_T1_10** | **B2** | ST141 | fliC_H6 | wzx_O50 wzy_O2 | 0 | 0 | 0 | 0 | 0 | 0 | 0 | 0 | 0 | 0 | 0 | 0 | 0 |
| **I7_T0_1** | **U** | ST1186 | fliC_H31 | wzx_O134 wzy_O134 wzy_O46 | 0 | 0 | 0 | 0 | 0 | 0 | 0 | 0 | 0 | 0 | 0 | 0 | 0 |
| **I7_T1_1** | **B2** | ST357 | fliC_H4 | wzx_O13/O129 wzx_O13/O129 | 0 | 0 | 0 | 0 | 0 | | | 0 | | | | 0 | |
| **I7_T2_13** | **B2** | ST141 | fliC_H6 | wzx_O50 wzy_O2 wzy_O2 | 0 | 0 | 0 | 0 | 0 | 0 | 0 | 0 | 0 | 0 | 0 | 0 | 0 |
| **I1_T0_22** | ***Enterobacter asburiae*** | NA | NA | NA | 0 | 0 | 0 | 0 | 0 | 0 | 0 | 0 | 0 | 0 | 0 | 0 | 0 |
| **I1_T2_14** | ***Enterococcus faecalis*** | NA | NA | NA | 0 | 0 | 0 | 0 | 0 | 0 | 0 | 0 | 0 | 0 | 0 | 0 | 0 |
| **I4_T1_11** | ***Klebsiella pneumoniae*** | NA | NA | NA | 0 | 0 | 0 | 0 | 0 | 0 | 0 | 0 | 0 | 0 | 0 | 0 | 0 |
| **I4_T2_1** | ***Citrobacter sp.*** | NA | NA | NA | 0 | 0 | 0 | 0 | 0 | 0 | 0 | 0 | 0 | 0 | 0 | 0 | 0 |
| **I6_T0_8** | ***Citrobacter freundii*** | NA | NA | NA | 0 | 0 | 0 | 0 | 0 | 0 | 0 | 0 | 0 | 0 | 0 | 0 | 0 |
| **I6_T2_17** | ***Enterococcus faecium*** | NA | NA | NA | 0 | 0 | 0 | 0 | 0 | 0 | 0 | 0 | 0 | 0 | 0 | 0 | 0 |
| **A4_T0_8** | ***Enterobacter cloacae*** | NA | NA | NA | 0 | 0 | 0 | 0 | 0 | 0 | 0 | 0 | 0 | 0 | 0 | 0 | 0 |
| **A4_T0_9** | ***Enterobacter cloacae*** | NA | NA | NA | 0 | 0 | 0 | 0 | 0 | 0 | 0 | 0 | 0 | 0 | 0 | 0 | 0 |
| **A6_T0_19** | ***Enterobacter cloacae*** | NA | NA | NA | 0 | 0 | 0 | 0 | 0 | 0 | 0 | 0 | 0 | 0 | 0 | 0 | 0 |
| **A7_T2_31** | ***Enterobacter cloacae*** | NA | NA | NA | 0 | 0 | 0 | 0 | 0 | 0 | 0 | 0 | 0 | 0 | 0 | 0 | 0 |

*Note no infectivity was observed against bacterial hosts at titres of $10^5$ PFUs/ml or less. * Phage infectivity was only observed for phage titres equal to or greater than $10^8$ PFU/s ml and values reported represent the comparably low efficiency of plaquing (EOP) relative to phage infectivity against indicator host *E. coli* MAC. Grey squares indicate turbid plaques, zero in white background = no infectivity.
